# Development of a wireless bioelectronic actuator for wound healing in a porcine model

**DOI:** 10.1101/2024.12.18.629068

**Authors:** Prabhat Baniya, Maryam Tebyani, Wan Shen Hee, Houpu Li, Narges Asefifeyzabadi, Kaelan Schorger, Hsin-ya Yang, Anthony Gallegos, Kan Zhu, Fan Lu, Gordon Keller, Celeste Franco, Koushik Devarajan, Alexie Barbee, Cristian Hernandez, Tiffany Nguyen, Cynthia Recendez, Elham Aslankoohi, Roslyn Rivkah Isseroff, Min Zhao, Marcella Gomez, Marco Rolandi, Mircea Teodorescu

## Abstract

Wireless bioelectronic actuators have been developed to deliver targeted treatments over multiple days while continuously monitoring delivery, thereby improving wound healing. Specifically, these devices can deliver charged biomolecules such as fluoxetine cations (Flx^+^) and electric field (EF) in freely moving pigs. Treatments can be controlled and monitored in real time via WiFi, with options for both user-specified delivery rates and durations, as well as automated closed-loop (CL) control. The devices are engineered to handle various failure scenarios that may arise in dynamic, real-world experiments—such as communication or power interruptions—ensuring that valuable experimental data is collected with minimal disruption. The ion pump features eight drug reservoirs and channels arranged around a 20 mm diameter-wound, with a central ground electrode (0 V). When voltages above 0 V are applied to the outer channels, currents flow from the reservoirs and channels into the wound, delivering Flx^+^ and/or EF depending on the reservoir solution. The device records applied voltages and currents locally to a microSD card at a high sampling rate, while simultaneously transmitting real-time measurements via a local WiFi network to a wound healing algorithm running on a nearby laptop. CL control of current/delivery rate is performed by an onboard microcontroller unit (MCU) and current-source microchips, based on instructions received from the wound healing algorithm. A graphical user interface (GUI) provides intuitive user control and real-time data visualization, with support for multiple devices. In vivo studies over seven days showed that Flx^+^-treated wounds had a 20% lower M1/M2 macrophage ratio and 41.67% greater re-epithelialization compared to controls (standard-of-care), demonstrating the actuator’s potential to enhance wound healing.

## Introduction

Wearable and implantable bioelectronic devices have applications in remote health monitoring^1^, wound healing^2^, diabetes, cancer, contraception, and the treatment of various other diseases^3^. Wound healing treatment remains one of the major medical challenges to date. Wound healing is an intricate physiological process that unfolds in four overlapping phases: hemostasis, inflammation, proliferation, and maturation^4–6^. Advancements in wound care include various types of smart dressings and patches that promote wound healing^7–9^ through thermo-responsive mechanisms for drug and antibiotic delivery, necessitating the use of microheaters and external power supplies^10^. Other traditional devices rely on wiring to bulky power supplies for applying exogenous electric fields with manual tuning of field strength to accelerate wound closure^11,12^, which limits their practicality for real-time treatment.

Self-powered and self-activated wearable piezoelectric^13^ and triboelectric nanogenerators for electrical stimulation (ES)^14,15^ and drug delivery^16^ have been demonstrated, but are limited to skin wound healing and offer little control over stimulation strength or drug release profiles. Electronic drug delivery (EDD) systems offer more precise spatiotemporal control of drug release rate, timing, and dosage, while reducing systemic side effects and improving patient compliance^17–19^ in wound healing applications^20^. In EDD systems, the rate and total dose of charged drug molecules are controlled by monitoring and adjusting the stimulation current amplitude and duration, respectively, enabling better drug penetration and sustained delivery into tissue compared to traditional oral and needle-based (hypodermic) methods^21–23^.

Moreover, wireless EDD devices provide ease of connectivity^24^ for real-time, continuous monitoring and remote control of drug dosage, allowing for localized, on-demand drug delivery^19,25–28^. Furthermore, wireless devices enable experimentation on a multitude of freely moving animals (each animal could carry multiple devices), providing more data and larger sample sizes per experiment, while minimally altering animal behavior compared to tethered systems^29^, thereby enabling better statistical analysis. Due to limited space at the wound site, small yet efficient antenna designs are essential for realizing reliable wireless links for bidirectional communication, despite variable antenna orientation, interference between multiple devices, and detuning in the presence of bodily tissues—particularly in ingestible bioelectronics^30^.

Notably, wearable wireless devices have also been demonstrated in vivo for accelerating wound healing. Most recently, a wireless bioelectronic system was demonstrated for antibiotic drug delivery (TCP-25 AMP) in combination with ES to treat infected chronic wounds in mouse models, while monitoring wound infection and inflammatory biomarkers such as temperature, pH, ammonium, glucose, lactate, and uric acid^31^. A wirelessly controlled micropump, in combination with a microneedle array, has been used for controlled release of VEGF with specific temporal profiles in diabetic mice^32^. This system achieves improved wound closure by utilizing microneedle arrays to transdermally reach deeper layers of the wound bed and improve the bioavailability and effectiveness of delivered drugs. Electronically controlled antibacterial cefazolin delivery through a closed-loop (CL) feedback system—using sensed temperature, pH, and uric acid levels at the wound site—has also been shown to treat infection and increase wound closure rates in rats^10^. The use of electrically stimulated hydrogel electrodes was demonstrated by^33^ to reduce wound size and increase dermal thickness in mice via CL control of stimulation in response to changes in wound impedance and temperature. Table 1 compares our work with other similar studies reported in the literature. Our fully integrated wireless bioelectronic device consists of polydimethylsiloxane (PDMS) reservoirs that hold charged drug molecules in solution and interface with the wound through hydrogel-filled capillaries^34^, while ensuring biocompatibility^35^. In this paper, we describe in detail the design of a printed circuit board (PCB) used to drive small electrical currents through the solution in the PDMS reservoirs and capillaries, as shown in Fig. 1a, to push charged drug molecules into the wound. The wireless bioelectronic device is validated and verified through in vivo experiments^34,36–38^.

**Table 1.** Comparison with other wireless wound-healing systems in animal models.

| Device | Drug delivered | Animal model | Actuation parameters | Wound healing mechanism |
| --- | --- | --- | --- | --- |
| Wireless bioelectronic actuator (this work) | Fluoxetine & EF | Pig | 0-4.7 V, 0-50 μA, 0-24 h DC/Pulsed DC | Reepithelialization M1/M2 |
| Wireless, stretchable bioelectronic system<sup>31</sup> | TCP-25 AMP & ES | Mice | 1 V, 1 mA, 10-20 min DC | Antibacterial |
| Wireless micropump/microneedle system<sup>32</sup> | VEGF | Diabetic mice | 3 V, 10 min | Vascularization |
| Wireless smart wound dressing<sup>10</sup> | Cefazolin | Rat | -0.5 V, 90 s DC | Antibacterial |
| Wireless, closed-loop, smart bandage<sup>33</sup> | ES | Mice | 0-2 V, 6 h, 13.56 MHz rectified AC | Proregenerative macrophage |
| Wireless, programmable, refillable hydrogel<sup>40</sup> | Metformin | Rat | 0-2 V, 30 min, 10 Hz AC/DC | Antibacterial Angiogenesis |
| Wireless, closed-loop, micropump microneedle array<sup>41</sup> | VEGF | Mice | 5.82-14.56 V, 1-7 mA DC | Tissue formation Collagen deposition |

**Figure 1.**
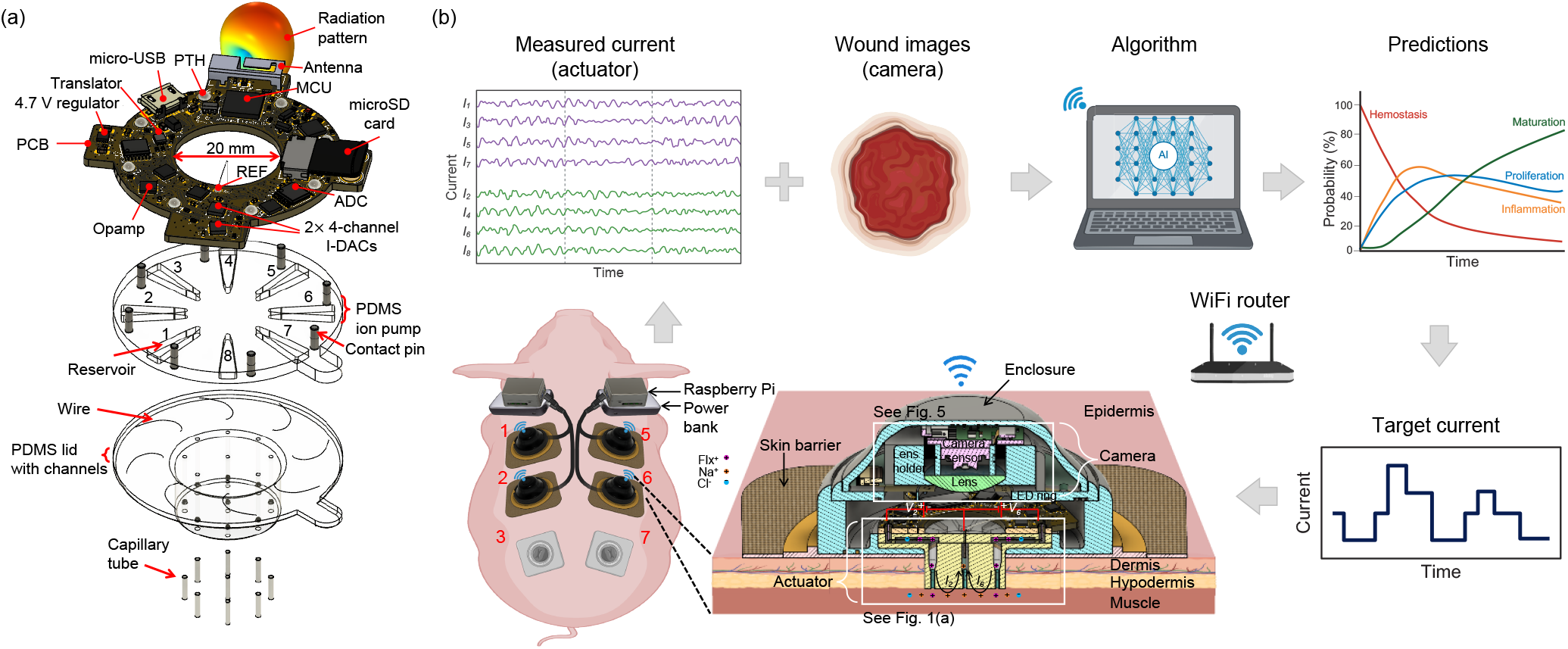
(a) Exploded view of the integrated PCB–PDMS wireless bioelectronic actuator. (b) Wireless bioelectronic devices are placed on the back of a pig to enable wireless communication with a laptop for real-time data acquisition and control of Flx^+^ and/or EF delivery for wound healing. The actuators measure the current passing through each wound, while cameras capture images of the wound surface. A wound healing algorithm running on the laptop processes these images to predict the wound healing stage. Based on these predictions, target current levels are set and transmitted to the actuator to control the current—and thereby the drug delivery rate. This cycle repeats until the target dose is achieved.

As illustrated in Fig. 1b, the electrical currents are monitored in real time and wirelessly through a graphical user interface (GUI) running on a nearby laptop. Users can also remotely control the devices in real time to start, stop, or adjust actuation as needed. The devices can additionally be placed in CL mode with the laptop to control the drug delivery rate based on target current levels set by a machine learning-based wound healing algorithm^39^.

## Results and discussion

### Experimental setup

Fig. 1b shows the experimental setup on a pig for multi-day Flx^+^ and/or EF delivery. The devices are placed on the back of a pig and communicate wirelessly with a nearby laptop through a local WiFi network that is set up using a commercial router.

The real-time data measured by the devices can also be remotely monitored and controlled through a 5G internet connectivity, as illustrated in Supplementary Fig. S1 online. Six wounds (left: 1, 2, 3; right: 5, 6, 7) were created on the pig. Wounds 1, 2, 5, and 6 were used as the treatment group and wounds 3 and 7 as the standard-of-care control group (no devices); see Supplementary Fig. S2 online. The actuated devices wirelessly receive commands from, and send measured voltages and currents to, a laptop where a wound healing algorithm^39^ determines the target current levels based on wound stage predictions obtained from wound images^42^.

### Design of wireless bioelectronic actuator

Fig. 1a shows the CAD model of the wireless bioelectronic actuator PCB with a WiFi/Bluetooth-capable microcontroller unit (MCU), microSD card, analog-to-digital converters (ADCs), and current-source digital-to-analog converters (I-DACs), with the PDMS device underneath. As shown in Fig. 1a, the PCB ring has an outer diameter of 43 mm and an inner diameter of 20 mm. There are 8 plated through-holes (PTHs), each 1.75 mm in diameter, where contact pins and electrode wires can be inserted to achieve mechanical and electrical integration with the PDMS device. The holes are located at a radial distance of 19 mm from the center. The simplified circuit diagram of the PCB is shown in Fig. 2a. The inter-integrated circuit (I^2^C) clock and data pins of the onboard MCU (Espressif ESP32-PICO-D4^43^) are connected to the respective pins of the I-DACs and ADCs to carry out instructions for setting the target currents (applied voltages are varied to achieve this) at the electrodes (actuation) and for reading the differential voltages at the current-sensing resistors (sensing), respectively. The PCB ring includes a micro-USB connector for powering the electronics. The connector is also used to flash the onboard MCU chip with the control program using a USB-A/C port on a laptop. As shown in Fig. 2a, the I-DAC outputs are connected to the contact pins in the PDMS, and the ADC measures the current *I*_*i*_ through each channel *i* by detecting the voltage drop across a high-precision (0.1%) current-sense resistor, *R*_*i*_ = 10 k*Ω*.

**Figure 2.**
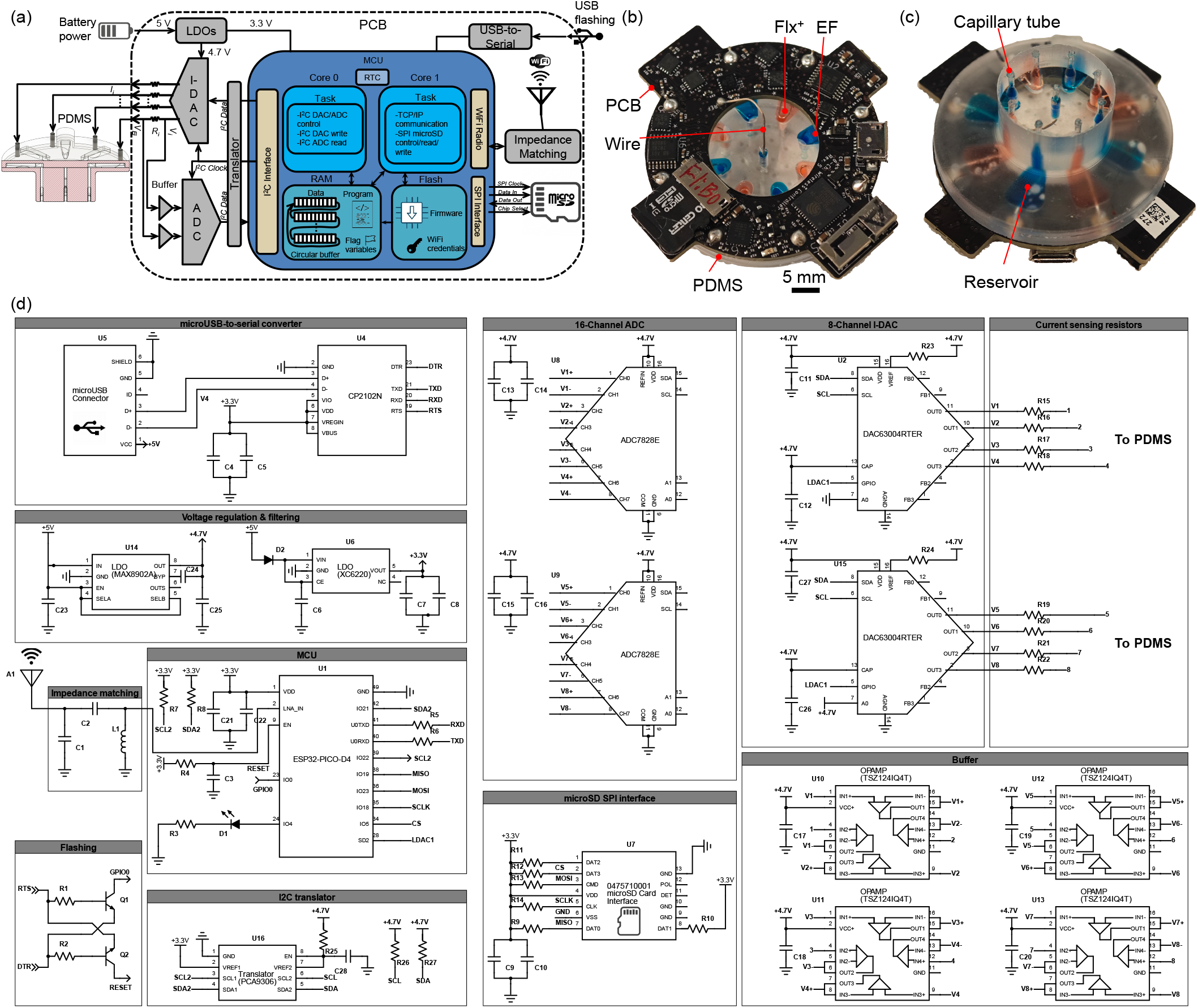
(a) Simplified circuit diagram of the wireless bioelectronic actuator PCB. (b) Top view of the PCB prototype with integrated PDMS. (c) Bottom view of the PCB prototype with integrated PDMS. d) Detailed circuit diagram of wireless bioelectronic actuator PCB showing the interconnection between the MCU, ADCs, I-DACs, opamps, microSD card, USB-to-serial converter, LDO regulators, power supply, flashing circuit, antenna, impedance matching network, SPI, and I^2^C interfaces.

The top and bottom views of the fabricated prototype integrated with PDMS are shown in Fig. 2b and c. The PDMS is filled with Steinberg solution^36,37^ (for EF) in the center and odd-numbered reservoirs and Flx^+^ solution^34^ in the even-numbered reservoirs. Before the PCB is integrated with the PDMS, each channel is separately and linearly calibrated in the 2 to 50 *µ*A range by measuring the currents passing through a 46.13 k*Ω* resistor (mid-scale value chosen although any resistor with < 84 k*Ω* resistance can be used so that I-DAC stays in the linear range without saturation) using a high-precision multimeter (see Supplementary Fig. S3 online). The resistors are removed after calibration and then integrated with the PDMS. The PCB has 8 PTHs for electrical and mechanical integration with the PDMS device. The integration is achieved with the help of 8 platinum (Pt)-coated pins that are inserted into the PDMS through-holes, as depicted in Fig. 1a. Each pin has a diameter and length of 1.6 mm and 4.8 mm, respectively. The pins are made of low-alloy steel but are Pt-coated to prevent corrosion. A conductive silver epoxy paste is manually coated on the bottom half before the pins are inserted into the PDMS through-holes. Therefore, each pin is electrically connected with the embedded wires (electrodes) in the PDMS holes. The 8 PTHs of the PCB mechanically align with the PDMS through-holes, allowing the PCB to be placed flush on the PDMS by guiding the PTHs over the pins. The protruding top portion of the pins is soldered to the pads of the PTHs to complete the PCB–PDMS integration, as shown in Fig. 1a. There is also a reference (REF) PTH, which serves as the electrical ground (0 V) of the PCB. A silver (Ag) wire is soldered to this PTH and inserted through the PDMS ion pump to establish an electrical connection with the center channel in the PDMS lid. After that, hydrogel-filled capillaries are manually inserted (using a push metal tool) in the 9 channel outlets at the bottom of the PDMS, as depicted in Fig. 1a. The capillaries provide a cation-selective electrical path between the reservoirs’ solution and the biological sample that is in contact with the device. Next, using a syringe, each reservoir is filled with either 10 mM Flx HCl solution^34^ or Steinberg solution^36,37^, depending on whether Flx^+^ or EF is to be delivered. Finally, the entire device is enclosed in a 3D-printed enclosure, as shown in Fig. 1b, to safeguard it from physical impacts resulting from animal movement.

The detailed circuit diagram in Fig. 2d shows the circuitry for the microUSB-to-serial converter for flashing, voltage regulation and filtering, impedance matching for the antenna, two 8-channel ADCs, two 4-channel I-DACs, current-sensing resistors, serial peripheral interface (SPI) for the microSD card, and operational amplifiers as buffers. An LED (D1) is also included, which connects to an MCU GPIO pin to indicate the WiFi connection status. It blinks when the MCU is attempting to connect to a WiFi network and turns off once the connection is established. An unregulated 5 V output from a power source (e.g., a power bank or battery) is regulated down to 3.3 V as the supply voltage for the MCU, and to 4.7 V as the supply voltage for the I-DACs and ADCs, using two separate low-dropout (LDO) regulators, as shown in Fig. 2d. Moreover, a Schottky diode (D2) is placed in series between the 5 V rail and the 3.3 V LDO input to provide reverse current protection for the MCU and USB port.

### I-DAC, ADC, and Translator

As shown in Fig. 2a, the actuation and sensing circuitry consists of I-DACs and ADCs, respectively. Both utilize a higher supply voltage of 4.7 V. Given that the 5 V output from most power sources is unregulated, it cannot be directly used to power the circuitry, since this would limit measurement accuracy. Therefore, an LDO regulator is incorporated to provide a regulated 4.7 V supply for the circuitry. The 4.7 V supply enables higher applied voltages and higher actuation currents than a 3.3 V supply. The I-DACs utilized are two Texas Instruments DAC63004, each having *N*_*dac*_ = 8-bit resolution with four outputs, and each can provide variable applied voltage *V*_*i*_ in the range from 0 to *V*_*CC*_ = 4.7 V at each of its outputs. Each output can provide a current *I*_*i*_ in the range from 0 to *I*_*max*_ = 50 *µ*A. As shown in Fig. 2a and d, each output is connected to a current-sense resistor, *R*_*i*_ = 10 k*Ω* ± 0.1%, which in turn is connected to an electrode (Ag or AgCl wires) in the PDMS. A 0.37 mm diameter Ag wire and an AgCl wire are used as the working electrode (WE) and reference electrode (RE), respectively. Each electrode can be set as either the WE (by setting *I*_*i*_ *>* 0 *µ*A) or the RE (by setting *V*_*i*_ = 0 V) for a specified duration. Additionally, each output can also be set to a high-impedance (Hi-Z) state, which electronically disconnects an I-DAC output from its electrode, resulting in a channel current *I*_*i*_ ≈ 0 *µ*A (open circuit). This feature allows the disabling of individual channels and provides better spatial control of actuation if/when needed. Moreover, the I-DACs enable rapid current control with negligible delay across all eight channels.

As shown in Fig. 2d, two eight-input ADCs are utilized to measure the voltage drop across each resistor *R*_*i*_. Operational amplifiers (op-amps) configured as voltage buffers are utilized at the ADC inputs to present a very high input impedance, which prevents current loading due to the ADCs and enables accurate measurement of actuation currents. A dedicated pair of inputs of the ADCs is connected across each resistor *R*_*i*_ to measure the voltage drop. The ADCs utilized are two Burr-Brown ADS7828, each with a *N*_*adc*_ = 12-bit resolution, which translates to 0.8 mV for voltage and 0.08 *µ*A for current measurements. Both the I-DAC and ADC support the I^2^C fast mode, in which the clock speed is set to 400 KHz, allowing the MCU to set or measure voltages at < 0.1 ms intervals. The I^2^C pins of the MCU, SCL and SDA, are pulled up externally by strong pull-up resistors (2 k*Ω*) to allow fast rise times (to meet setup and hold time requirements) needed for the fast mode. The ADCs sample the analog applied *V*_*i*_ and electrode 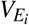 voltages across each resistor *R*_*i*_ and convert them into 12-bit integers for each channel. A burst of sixteen samples is taken at each input, spaced 0.5 ms apart. The MCU then finds the average of these samples to reduce noise. The sampling interval between the averages is 100 ms. The MCU stores these point-averaged *N*_*adc*_ = 12-bit integer codes *A*_*i*_ and 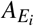, for applied voltage and electrode voltage, respectively, on the microSD card via the SPI bus for each channel *i*. The average applied *V*_*i*_ and electrode 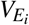 voltages at each channel *i* can be found as

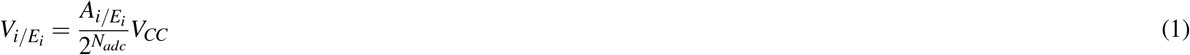

The voltage drop, 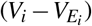, across resistor *R*_*i*_ can then be used to find the current flowing at each channel *i* as

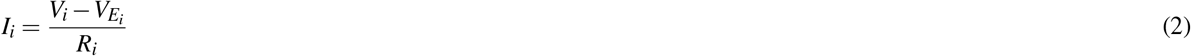

Eqs. (1) and (2) allow calculation of voltage and current at each channel when the ADC and I-DAC codes are not calibrated and represent the ideal voltage and current transfer functions of the device. When both ADC and I-DAC codes are calibrated using the two-point calibration method, the voltage and current at each channel are calculated using Eqs. (2) and (12), respectively, resulting in more accurate measurements (since the noise is calibrated out) and representing the calibrated transfer functions of the device.

The ideal and calibrated voltage transfer functions of the ADC for channel 1, using a load resistance of *R*_1_ = 80.4 k*Ω* (near full load resistance chosen so that the input voltage can swing the entire 0-4.7 V linear range without saturation), are shown in Fig. 3a. The ideal and calibrated current transfer functions of the I-DAC for channel 1 are shown in Fig. 3b. These are obtained from Supplementary Eqs. (1) [ideal ADC], (2) [calibrated ADC], (5) [ideal I-DAC], and (6) [calibrated I-DAC], as the I-DAC output code *D*_*i*_ is varied from 0 to 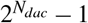, i.e., 1 to 50 *µ*A current output. The absolute errors are also included and are calculated as the difference between the device and true voltages/currents. At each point (2 *µ*A steps), true values of voltages and currents at the channels are recorded with a multimeter (see Supplementary Fig. S3 online), whereas the device values (averaged from 200 samples = 20 s for each step) are recorded by the device itself. The absolute voltage error (mV) increases linearly with higher input voltage (and by extension, input current), and the I-DAC absolute current error (*µ*A) increases linearly with higher input code up to near full-scale (4.7 V), after which both drop sharply due to saturation (non-linearity) of the I-DAC. The multimeter measurements (labeled “measured”) lie closer to the calibrated transfer function curves, which is to be expected since they are recorded after the device is calibrated and serve as an additional cross-check. The measured curves somewhat bend and don’t increase proportionally near the upper limit, in contrast to a straight diagonal line (ideal/calibrated response). The deviation between the ideal and calibrated voltage transfer function for the ADC is negligible, as evident in Fig. 3a, but some deviation for the I-DAC is noticeable in Fig. 3b, particularly as a small positive full-scale error (offset plus gain error). Other channels also show similar performance, as shown in Supplementary Figs. S4 and S5 online. The maximum absolute voltage error is 70 mV, and the maximum absolute current error is 1.25 *µ*A, both of which occur at channel 3 when the input voltage is 4.2 V.

**Figure 3.**
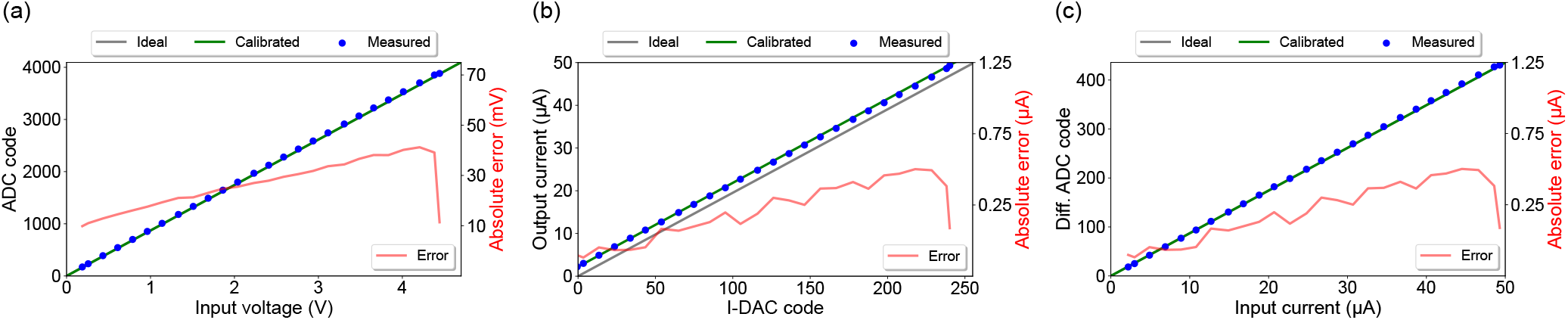
Channel 1 transfer functions. (a) ADC transfer function for voltage measurements. (b) I-DAC transfer function for current output. (c) ADC transfer function for current measurements.

The ideal and calibrated current transfer functions of the ADC for channel 1, again using a load resistance of *R*_1_ = 80.4 k*Ω*, along with the absolute error between the device and true currents, are shown in Fig. 3c, which are obtained from Supplementary Eqs. (9) and (11), respectively. True current values in 2 *µ*A increments are recorded using a multimeter placed in series with the load resistance. The multimeter measurements (labeled “measured”) closely follow the calibrated current transfer function curve, indicating high current measurement accuracy within the full-scale 50 *µ*A range after calibration. The ADC’s absolute current error (in *µ*A) also increases linearly with input current, but drops sharply near full-scale due to saturation-induced non-linearity. Other channels also show similar performance as shown in Supplementary Fig. S6 online.

As illustrated in Fig. 2a and d, the MCU and microSD card are powered by a 3.3 V rail, as they do not support higher voltages. For I^2^C communication, a 3.3 V to 4.7 V level translator chip is employed to enable communication between the 3.3 V MCU and the 4.7 V ADCs/I-DACs. The translator resolves logic-level mismatches by mediating the SCL (clock) and SDA (data) lines between the MCU and the peripheral devices.

Table 2 summarizes the main controller specifications. I^2^C communication between the MCU (master) and I-DACs/ADCs (slaves) proceeds as follows. To start communication, the master sends a START condition on the I^2^C bus (switching SDA line from HIGH to LOW followed by switching SCL line from HIGH to LOW) (see Fig. 2a and d). The MCU then transmits a 7-bit slave address along with a read (HIGH) or write (LOW) bit—together forming the address byte—on the SDA line. All the slaves read each bit serially. Each bit is read when the SCL is HIGH, requiring 8 clock cycles. The slave with the matching address acknowledges by pulling SDA LOW on the 9th clock cycle. After that, the MCU typically sends a control byte to configure the slave (e.g., select an ADC/I-DAC channel to read/write, power up/down channels, etc.), which is acknowledged by the slave. Then, one or more data bytes are transferred between the MCU and the slave (e.g., read a voltage measurement from the ADC, set a voltage or current at the I-DAC output). After each byte is transferred, the receiver acknowledges the sender. Finally, to stop communication, the master sends a STOP condition (switching SCL line from LOW to HIGH followed by switching SDA line from LOW to HIGH).

**Table 2.** Specifications of wireless biolectronic actuator.

| Parameter | Specifications |
| --- | --- |
| Channels | 2× 4-actuation (I-DACs), 2× 8-sensing (ADCs) |
| Applied voltage $V_i$ | 0 to 4.7 V |
| Channel current $I_i$ | 0 to 50 μA |
| Actuation resolution | 8-bit I-DACs (0.2 μA) |
| Sensing resolution | 12-bit ADCs (1.1 mV) |
| Averaging factor | 16 (0.5 ms apart) |
| Sampling period between averages | 100 ms |
| Power management | Active/light sleep/standby |
| Typical delivery duration | 24 h per day (up to 7 d) |
| microSD card size | 4 to 32 GBytes |
| Current sensing resistors $R_i$ | 10 kΩ ±0.1% |

### GUI and laptop software

As illustrated in Fig. 1b, the wireless bioelectronic actuator is remotely operated via a GUI on a laptop connected to a local WiFi network established using a commercial router. To verify that the device is functioning correctly, real-time current and voltage measurements are first validated by setting a target current across a known resistor at all eight outer and center channels, respectively (see Supplementary Fig. S3 online).

The GUI and backend data processing software are implemented in Python using the Spyder (Anaconda3) IDE. The user flow diagram for the GUI, laptop software, and device firmware (instruction flow) is shown in Fig. 4. The GUI includes push buttons for scanning the WiFi network to detect all connected devices (Find Devices), a drop-down menu to select a device from among those found, loading the actuation table (Load), showing the actuation table, enabling CL control (Close Loop), calibrating the I-DACs and ADCs (Calibrate), real-time viewing of measured voltages and currents (View Readings), displaying SD card data stored on the device (Show SD Card Data), transferring files from the device to the laptop (Copy), deleting files on the device (Delete), and starting (Start) or stopping (Stop) the actuation (see Supplementary Fig. S7 online).

**Figure 4.**
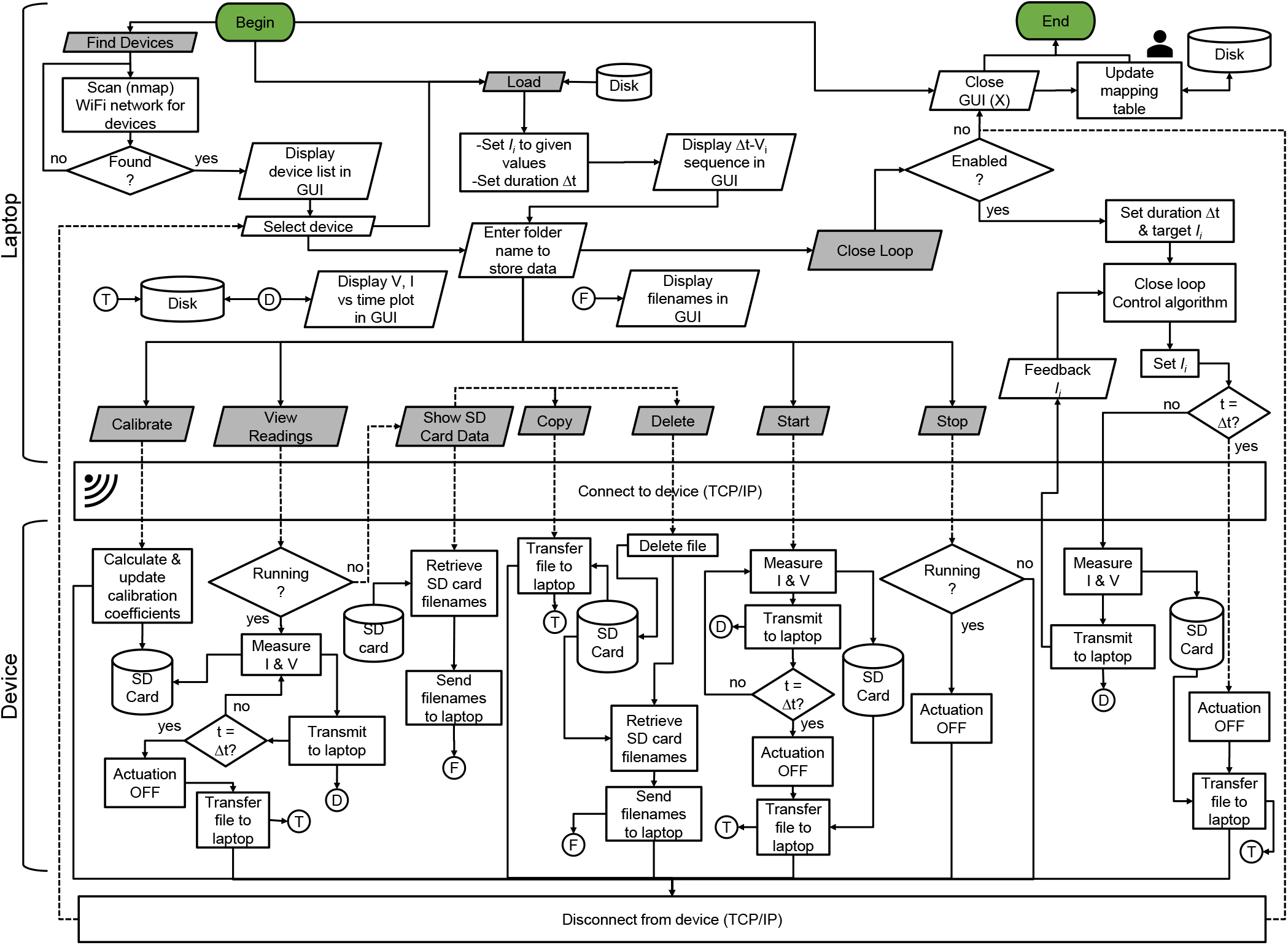
Flowchart for data acquisition and control, showing the communication between a laptop and a wireless bioelectronic actuator. The GUI running on the laptop includes several buttons, represented by gray parallelograms. Each button press typically establishes a TCP/IP connection with a selected device via the local WiFi network. Depending on the button pressed, the device performs specific tasks such as calibration, viewing measurements, showing SD card data, copying files, deleting files, starting open-loop actuation, stopping actuation, or enabling CL control.

When powered up, the device automatically connects to the network, and the router assigns it a unique IP address. The laptop is also connected to the same network. When the Find Devices button is pressed in the GUI, the software scans the network to detect all connected devices, each of which is uniquely identified by its media access control (MAC) and IP address. When a user initiates a specific action on a selected device (e.g., by pressing the Start button), the laptop first establishes a TCP/IP connection with the device and sends a high-level command indicating the action to be performed, as depicted in Fig. 4. On the device side, each command is uniquely identified, and the appropriate sequence of tasks is executed by the firmware.

For example, during actuation, a fixed-duration sequence of target currents is first loaded from a .csv file (actuation table) by pressing the Load button and selecting the file on the laptop. Next, pressing the Start button instructs the firmware to apply the specified target currents at the channels, measure the resulting voltages and currents, and wirelessly transmit the data in real time to the laptop for immediate viewing and/or CL control. Simultaneously, for redundancy and reliability, the data is also logged locally on the device’s microSD card at a higher sampling rate, leveraging its lower latency compared to WiFi transmission. After the actuation completes, the SD card data is automatically transferred to the laptop. The sequence of tasks performed by the laptop and device for all GUI functions is detailed in the flowchart shown in Fig. 4.

### Device firmware and MCU flashing

When a button is pressed on the GUI, high-level commands are sent wirelessly to the device, where the firmware breaks them into a sequence of low-level tasks for the MCU to execute, as depicted in Fig. 4. The device firmware is a control program written in C++ using the Arduino IDE environment. Once the device completes the action, it disconnects from the laptop and enters standby mode so that it can accept further commands from the laptop. As shown in Fig. 2a and d, the PCB houses a Universal Serial Bus (USB)-to-Universal Asynchronous Receiver and Transmitter (UART) bridge called CP2102N (an IC from Silicon Labs) that allows direct connection to a laptop’s USB port for flashing the MCU with custom firmware and debugging the firmware, if needed.

The main tasks the firmware performs are TCP/IP communication, data acquisition (through the ADCs), control (via the I-DACs), and reading from/writing to the microSD card, distributed between the two cores of the MCU, as depicted in Fig. 2a.

The microUSB connector on the PCB is used as an interface for flashing and powering purposes via a USB-A cable connected to a laptop or power source. The microUSB connector’s D+ and D-data pins are connected to the corresponding pins of the UART bridge for data transfer, as shown in Fig. 2d. The bridge itself is connected to the MCU for transmitting and receiving UART data through the TxD and RxD pins^43^. The process of flashing the PCB takes less than 1 min due to the 3 Mbps transfer rates supported by the UART bridge. The MCU is put into flashing mode by the UART bridge through the use of RTS and DTR pins, which are connected to two transistors that control the state of the MCU pins IO0 and EN, as shown in Fig. 2d. By putting the IO0 and EN pins through a set sequence of logic HIGH and LOW states, the UART instructs the MCU to initiate flashing mode for downloading firmware. The firmware is then transferred to the flash memory of the MCU via the RxD pin. All of these low-level operations are handled internally by the Arduino IDE during the flashing process. Print statements are used in the firmware to send device status updates to the UART bridge via the TxD pin, and then to the laptop through the microUSB connector for display/debugging on the Arduino IDE’s serial monitor. The firmware can be updated easily by re-flashing the MCU’s memory.

The firmware takes advantage of multicore processing. The MCU on the device has two cores: 0 and 1, as depicted in Fig. 2a. Core 0 is dedicated to sampling data from the ADCs and changing the I-DAC output voltage, while Core 1 is dedicated to handling TCP/IP commands for WiFi communication with the laptop and reading/writing to the microSD card. During actuation, the data from the ADCs come at a faster rate than can be transmitted wirelessly. Core 0 stores the measured ADC data into a sequence of allocated memory buffers, while Core 1 sends the data in a first-in, first-out (FIFO) fashion to the laptop while also recording it on the microSD card. This allows the system to record data at higher sampling rates than would otherwise be possible. The number of buffers and buffer size must be high enough to prevent overfill and data loss, and is determined by the worst-case latency of WiFi transmission, which is on the order of a few seconds. As illustrated in Fig. 2a, the buffers are referenced in a circular fashion for efficient utilization of memory. Intercore communication is handled through flag variables in the integrated 4-MB SPI flash of the MCU. The data is continuously sampled, only pausing momentarily (< 1 s) to avoid race conditions when a new CL command is received and the target currents and applied voltages must be changed.

### microSD card interface

The SPI communication protocol is used for data transmission between the MCU (master) and the microSD card (slave), as illustrated in Fig. 2a and d. The four default SPI pins of the MCU — MOSI (23), MISO (19), SCLK (18), and CS (5) — are connected to the CMD, DAT0, CLK, and DAT3 pins, respectively, of the microSD card connector to provide full-duplex communication. SPI Mode 0 is used for data transfer. This means that the SCLK (clock) signal stays LOW when there is no data transmission (i.e., when CS is HIGH), and data is simultaneously received (on the MISO pin) on the rising edge and transmitted (on the MOSI pin) on the falling edge of the clock by the MCU.

As indicated in Fig. 2d, the SPI pins are pulled up by 10 k*Ω* resistors to provide faster rise times required by high-speed microSD cards, which cannot be achieved using the weak internal pull-up resistors (45 k*Ω*) of the MCU. Pull-up resistors of 10 k*Ω* are also used on the SD card pins DAT0–DAT3, even though they are not connected to the MCU (since SPI mode is being used), to prevent the card from entering an incorrect state.

### Antenna and impedance matching

As shown in Fig. 1a, a 3D metal antenna is used for transmitting and receiving 2.45 GHz radio signals for WiFi communication. The antenna is tuned to this frequency using an impedance-matching network, which is connected to the LNA pin of the MCU, as illustrated in Fig. 2d. The *π*-matching network maximizes power transfer between the LNA and the antenna at 2.45 GHz. The PCB board material, thickness, trace width/length, and the proximity of other electronic components dictate the values of capacitors *C*_1_, *C*_2_, and inductor *L*_1_ required for adjusting the resonant frequency of the antenna.

Based on the manufacturer’s recommendation, the following values were utilized: *C*_1_ = 2.7 pF, *C*_2_ = 4.7 pF, and *L*_1_ = 1.6 nH^43^. These values are used to cancel out the parasitic reactance (capacitance and inductance) associated with transmission lines routed from the LNA to the antenna input. The antenna is surface-mounted at the edge of the board to minimize resonant frequency shifts and interference from nearby circuits/components.

The PCB comprising the matching network and the antenna was simulated in COMSOL using the finite element method. The reflection coefficient magnitude |*S*_11_| at the matching network input, with the antenna as its load, is –15.3 dB at 2.45 GHz, which indicates that 97% of the power provided by the LNA is accepted by the antenna and the network. The maximum gain of the antenna is 1.2 dBi. Fig. 1a shows the 3D gain pattern of the antenna and how it aligns with the structure of the antenna itself. Supplementary Fig. S8 online shows the 3D, horizontal, and vertical plane radiation patterns of the antenna.

### Closed-loop (CL) control of current

Fig. 5 shows a simple high-level diagram of how the wireless bioelectronic device typically operates in conjunction with a wound healing algorithm^39^ running on a laptop to ensure CL control of current for Flx^+^ delivery. When a positive voltage *V*_*i*_ is applied between the WE at the reservoir and the RE at the center of the PDMS ion pump, drug cations (e.g., Flx^+^) are pushed through cation-selective hydrogel-filled capillaries from the reservoir into the wound bed in exchange for endogenous sodium cations (Na^+^) at the RE. Flx^+^ is delivered from even channels 2, 4, 6, and 8 until a target dose is reached. The center and odd channels are filled with Steinberg solution (for EF) and grounded (0 V) during Flx^+^ delivery to reduce delivery resistance and improve delivery efficiency. The currents from even channels are time-integrated individually, scaled by delivery efficiency, and added together to determine the total dose delivered in real-time. The device can also deliver EF from odd channels 1, 3, 5, and 7, and this involves a similar CL control of current until a target duration is reached (integration and dose calculation steps are skipped in Fig. 5). Even channels are all open-circuited (Hi-Z, 0 *µA*) so that Flx^+^ is not delivered. This also ensures that the direction of the applied EF is radially inward, since the center channel is always grounded (0 V). The target current range is also wider and higher for EF delivery. Table 3 lists the target current ranges for Flx^+^ and EF delivery. The CL begins by setting the target current levels based on wound stage probabilities calculated from wound images captured by the cameras^42^ and wirelessly transmitted to the laptop. The target currents are calculated by the algorithm on the laptop and then wirelessly transmitted to the device. To maintain target levels, the onboard I-DACs on the device rapidly adjust (increase or decrease) the applied voltages at each channel without noticeable delays. The current can change due to the dynamic nature of wound and contact resistances, which are manifestations of bodily fluids and animal movement, respectively. However, such changes are rapidly compensated for by the I-DACs through rapid hardware-based feedback, which involves adjusting the applied voltage quickly to minimize current fluctuations. This can be seen in Fig. 5, where the applied voltage (red) is adjusted to keep the current (blue) at the target level (black). Moreover, the device firmware has been optimized to reduce communication latency by efficiently multitasking between data acquisition/control and handling TCP/IP commands from the laptop. This also significantly reduces data acquisition gaps and control action delay, enabling the device to quickly respond when the target current changes. The currents are pulsed (which is why they appear filled) at a 92.3% duty cycle (target value for 60 s, 0 *µ*A for 5 s) to allow electrodes to charge and discharge until the target dose of 0.025 mg is reached, after which all channels are grounded (0 V) to stop delivery. The CL control operation can run for an entire day or until the target dose/duration is reached, whichever is earlier, as the camera captures new images of the wound and updates the probabilities of the wound stage.

**Table 3.** Target currents and voltages for Flx^+^ and EF delivery.

| Delivery | Ch. 1 | Ch. 2 | Ch. 3 | Ch. 4 | Ch. 5 | Ch. 6 | Ch. 7 | Ch. 8 |
| --- | --- | --- | --- | --- | --- | --- | --- | --- |
| EF | 4.4-50 $\mu\text{A}$ | 0 $\mu\text{A}$ | 4.4-50 $\mu\text{A}$ | 0 $\mu\text{A}$ | 4.4-50 $\mu\text{A}$ | 0 $\mu\text{A}$ | 4.4-50 $\mu\text{A}$ | 0 $\mu\text{A}$ |
| $\text{Flx}^+$ | 0 V | 2.45-5 $\mu\text{A}$ | 0 V | 2.45-5 $\mu\text{A}$ | 0 V | 2.45-5 $\mu\text{A}$ | 0 V | 2.45-5 $\mu\text{A}$ |
| Stop | 0 V | 0 V | 0 V | 0 V | 0 V | 0 V | 0 V | 0 V |

**Figure 5.**
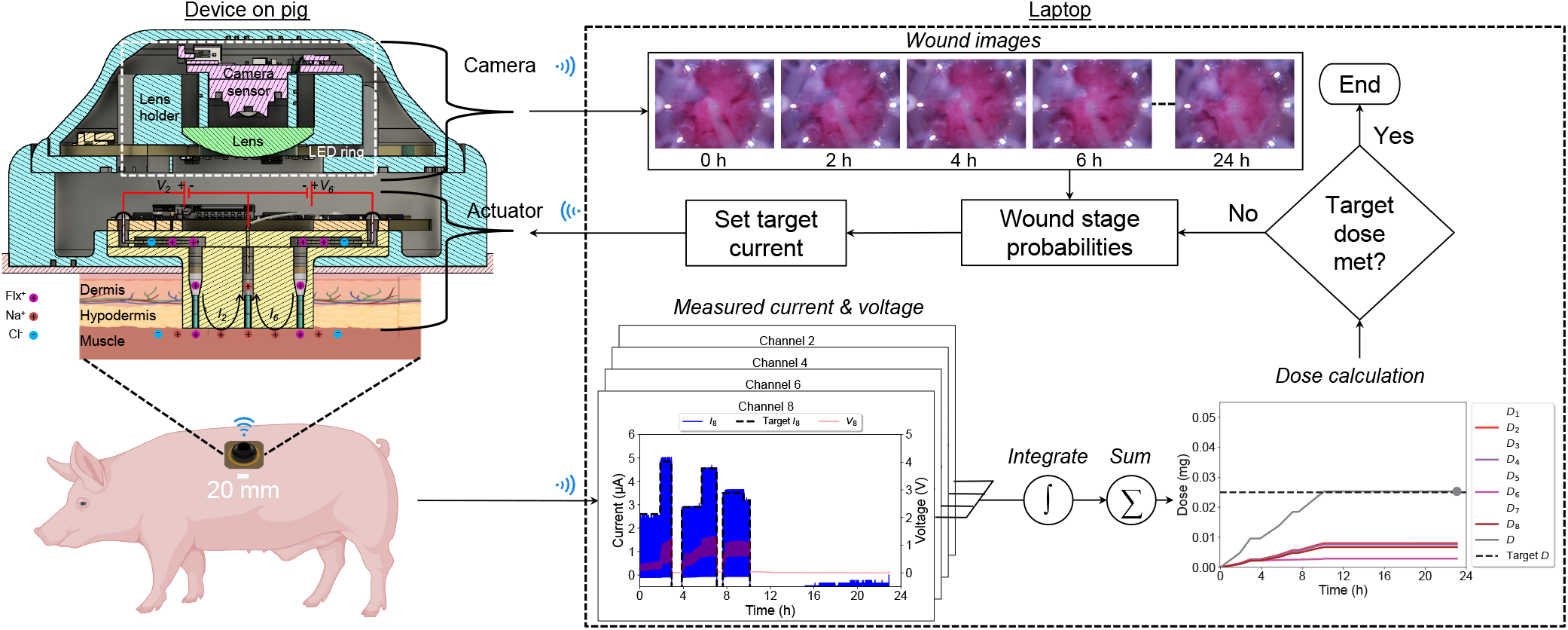
CL control of current in the wireless bioelectronic device using onboard I-DACs with pulsed current. The target currents are determined by the wound healing algorithm running on a laptop, based on wound stage probabilities calculated from wound images captured by the cameras.

### Power consumption

Table 4 shows the different operating modes of the device and the typical current consumption in each mode. Just before the start and immediately after the end of an experiment, the devices are typically in standby mode, in which an actuation can be started. Normally, during Flx^+^ or EF delivery, the device is in actuation (transmission) mode, and data is transmitted in real-time to a laptop. Sometimes, the laptop can be disconnected from the device for power-saving purposes while allowing the devices to continue delivering. This mode is called actuation (no transmission), but CL control and real-time data viewing features are lost. The devices also take advantage of sleep mode to reduce overall power consumption when they are not actuating, especially during long in vivo experiments. Since each device can generally reach the target dose within 6 to 12 h, the device can be put into sleep mode for the rest of the day to save power. During the experiment, the power banks connected to the devices can only be replaced once a day due to the limitations and protocols of animal handling. The use of sleep mode provides enough power savings to allow multiple devices to run and actuate on their respective wounds each day.

**Table 4.** Current consumption by the wireless bioelectronic actuator across different operating modes.

| Operating mode | Current (mA) | Comments |
| --- | --- | --- |
| Sleep | < 10 | No actuation, WiFi disabled, can be woken up after a preprogrammed time to put into standby mode |
| Standby | ≈ 50 | No actuation, WiFi connected, ready to accept commands |
| Actuation<br>(No transmission) | 70-150 | Data recorded on SD card, no transmission to laptop |
| Actuation<br>(Transmission) | 70-170 | Data recorded on SD card and real-time transmission to laptop via WiFi |

### Failure resiliency

Software and firmware improvements have been implemented on the laptop and device to handle occasional communication and power loss scenarios. When communication failures occur, network exceptions are raised in the software and handled on the laptop by continuously attempting to reestablish the connection with the device. Moreover, a resume feature has been incorporated into the firmware to resume interrupted actuation resulting from a power failure. Since the last recorded state (timestamp, duration, and target currents) of the actuation prior to power loss is stored on the microSD card, this information is used to continue actuation, and shortly after, the laptop can reestablish the connection with the device to resume CL control.

### Experimental results

As shown in Fig. 1b, the wireless bioelectronic devices were used on pigs to deliver EF and/or Flx^+^ for 7 days. After the experiment, both the treated and control wounds were analyzed and compared in terms of wound healing metrics such as the M1/M2 macrophage ratio and re-epithelialization. For Flx^+^ delivery on a wound, the current level dictates the delivery rate, which, in combination with the actuation duration, dictates the total dose delivered. The actuation duration for delivery was set to 23 h per day to reach a target dose of 0.025 mg per wound, but the delivered dose may reach the target earlier, as shown in Fig. 5, in which case the channels are grounded (0 V) to stop the delivery for the remainder of the period. The remaining 1 h allows for power bank replacement and animal care (food and water) in preparation for actuation over the next 23 h. For EF delivery, the current level dictates the strength of the applied EF, and the current range is set by the algorithm based on the wound stage probabilities. The actuation duration for EF delivery is also limited to 23 h per day for the same reason as Flx^+^ delivery.

Some representative experimental results from the device, showing 7-day EF (Day 0–1) and Flx^+^ (Day 1–6) delivery in a pig wound, are presented in Fig. 6. The measured currents and voltages on Day 1 at all eight channels are shown in Fig. 6a. The temporal progression of Flx^+^ dose per channel (*D*_*i*_) over a period of 7 days is shown in Fig. 6b. The spatial distribution of *D*_*i*_ for each day is shown in Fig. 6c. CL control of current is performed to achieve and maintain specific target levels, as shown in Fig. 6a for Day 1 actuation, when the treatment is switched from EF to Flx^+^ at the 6-hour mark. On Day 0 and before this mark, EF is delivered from the odd channels (top row), as seen by the pulsed nature of the current (blue) at channels 1, 3, 5, and 7, where the top envelope tries to follow the changing target current (black), whereas the even channels 2, 4, 6, and 8 (bottom row) are set to 0 *µ*A to disable Flx^+^ delivery. After this mark, Flx^+^ is delivered from the even channels (bottom row), whereas the odd channels are set to 0 V (red) to stop EF delivery, following the scheme of Table 3. On Day 0, as shown in Fig. 6b, only EF is delivered and Flx^+^ is not delivered at all, based on the recommendations made by the wound healing algorithm. The decision on when to switch from EF to Flx^+^ is also made by the algorithm.

**Figure 6.**
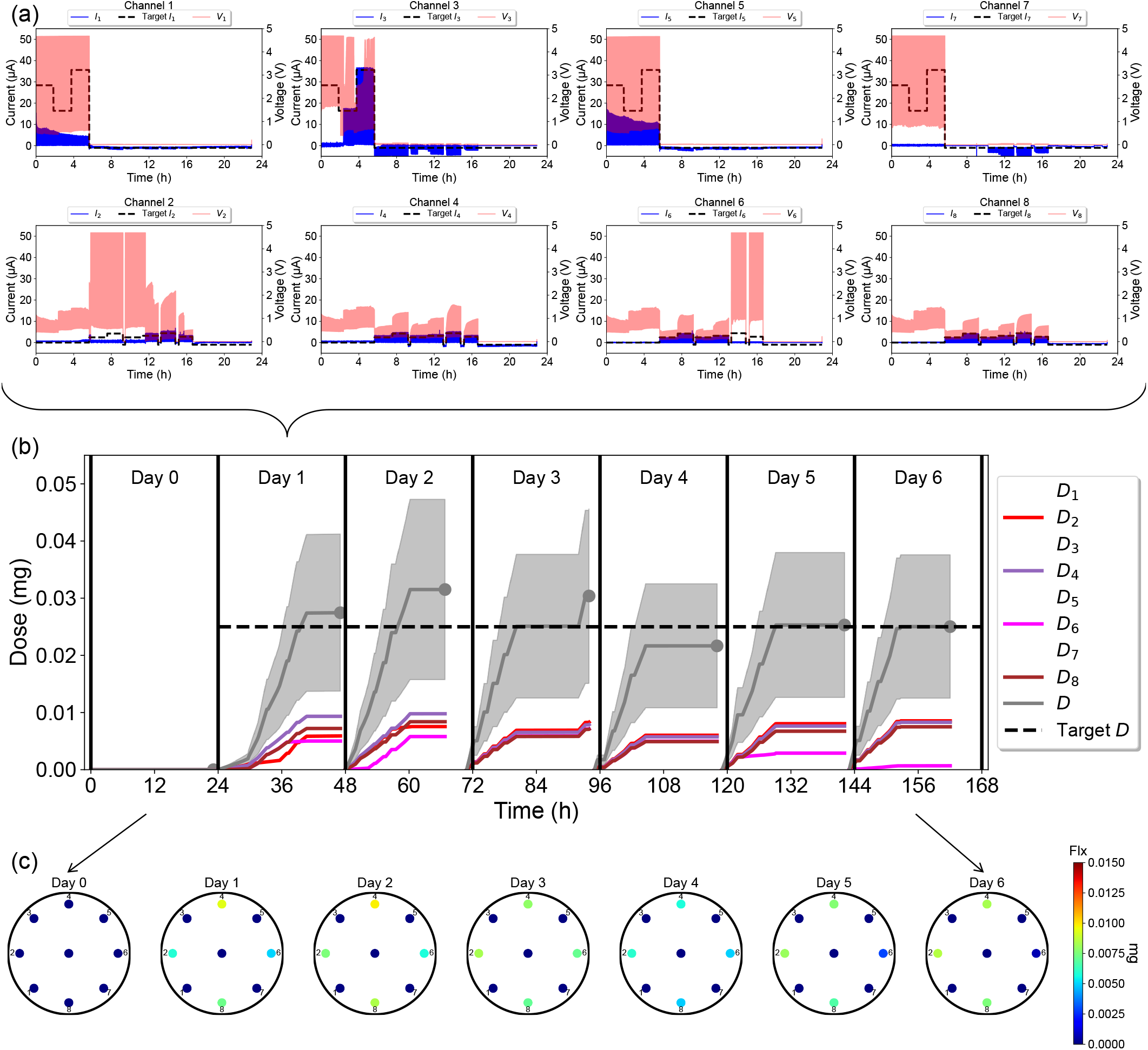
Experimental data from 7-day EF/Flx^+^ delivery on a wound. (a) Measured applied voltages and currents at the 8 channels on Day 1. (b) Temporal variation of accumulated per-channel and total dose during the 23 h delivery window on each day. All data presented as mean ± s.d. (c) Spatial variation of accumulated per-channel and total dose delivered at the end of the 23 h delivery window on each day.

### Fluoxetine dose calculation

For Flx^+^ delivery, the four even-channel reservoirs in the PDMS ion pump shown in Fig. 1a are filled with a 10 mM Flx HCl solution^34^, and the corresponding capillaries are loaded with Flx^+^ cations. The currents measured by the wireless bioelectronic actuator are directly proportional to the rate of Flx^+^ delivery. To quantify the total Flx^+^ dose delivered, the current at each even channel is cumulatively integrated and summed, as diagrammatically indicated in Fig. 5. The accumulated charge exchange, *Q*_*i*_, at channel *i* at time *t* can be calculated as

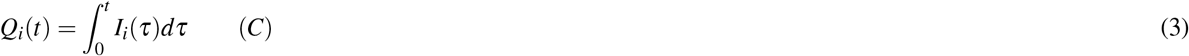

The total accumulated charge exchange, *Q* (in coulombs), is the sum of the charge exchanges at all even channels, from *i* = 2 to 8, and is given by

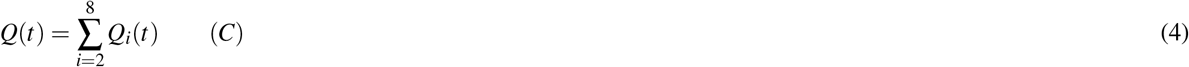

For Flx^+^ treatment, the accumulated dose *D*_*i*_ at channel *i* at time *t* can be calculated by integrating only the positive currents 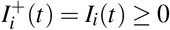 as follows:

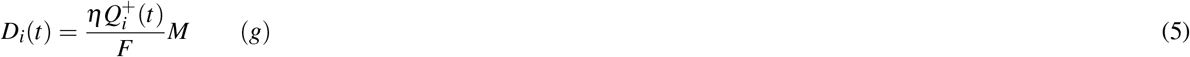

where *η* = 2.2 ± 1.1% [mean *±* standard deviation (s.d.)] is the Flx^+^ delivery efficiency of the device, based on HPLC calibration in vitro^34^; *M* = 309.33 g/mol is the molar mass of Flx; *F* = 96485.3321 C/mol is the Faraday’s constant; and 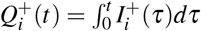.

The reason why only contributions of positive currents are integrated is because Flx^+^ delivery is assumed to occur only when the current is positive (i.e., Flx^+^ cations flow from the reservoirs through the cation-selective hydrogel to the wound bed and then diffuse away into the bloodstream), as illustrated in Fig. 5. When the current is negative, endogenous cations such as Na^+^ carry the current back to the reservoirs to maintain charge balance and do not reverse or cancel out Flx^+^ delivery. Hence, the positive and negative currents correspond to the exchange of different types of ions—Flx^+^ and endogenous cations, respectively.

The total accumulated dose, *D* (in grams, g), can be calculated by summing the accumulated channel doses, *D*_*i*_, at all even channels *i* = 2 to 8, as follows:

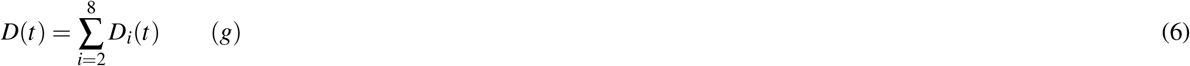

Using Eqs. 5 and 6, the temporal variation of accumulated channel dose *D*_*i*_ and total dose *D* from the measured currents (shown in Fig. 6a) was calculated for Day 1, as shown in Fig. 6b. The spatial distribution of channel dose *D*_*i*_ at the wound edges and center at the end of Day 1 is illustrated in Fig. 6c. The dose delivery is non-uniform across the wound; for example, see Day 1, *D*_4_ *> D*_8_ *> D*_2_ *> D*_6_, even though the total dose *D* ≈ 0.025 mg is close to the Day 1 target.

This non-uniformity is attributed to variation in current at each channel, caused by differences in contact quality, channel geometry, and wound resistance. If the resistance at a particular site is too high (e.g., due to poor contact or faulty capillaries), the maximum voltage (i.e., *V*_*CC*_ = 4.7 V supply) may be insufficient to achieve the target current, resulting in reduced delivery. The dose delivery enabled by the bioelectronic device is gradual, which helps improve bioavailability, unlike oral or topical drug administration.

The start of actuation is marked by a vertical line in Fig. 6b. Actuation typically begins between 12:00 PM and 2:00 PM and continues for 23 hours. The total accumulated dose *D* (represented by a gray dot) varies slightly from day to day, despite a constant target dose. This small deviation is most likely due to numerical integration errors accumulating over time during dose calculation.

In Fig. 6b, the slope of each *D*_*i*_ curve reflects the delivery rate at that channel and serves as an indicator of individual channel performance. A higher slope corresponds to a higher delivery rate, while a flat *D*_*i*_ (i.e., zero slope) indicates zero delivery. For instance, on Day 2 after the 60-hour mark, *D*_2_, *D*_4_, *D*_6_, and *D*_8_ exhibit near-zero slopes, suggesting near-zero or negative currents, and thus, no further Flx^+^ delivery, as the target dose had already been reached. Conversely, before the 60-hour mark, those same channels have positive slopes, indicating active Flx^+^ delivery.

The central channel exhibits negative currents (not shown), as it serves as the return path for the channel currents *I*_*i*_ and does not contribute to delivery, as evidenced by the blue dots (0 mg) at the wound center in all dose maps shown in Fig. 6c. On Day 0, no Flx^+^ is delivered from any channel. From the temporal and spatial dose profiles in Fig. 6b and c, it is evident that all four delivery channels function on Days 1–4, while only three are functional on Days 5–6.

The actual fluoxetine concentration in the wound tissue correlates with the cumulative dose (see Supplementary Fig. S9a online), but was not detected in plasma (see Supplementary Fig. S9b online), suggesting localized, rather than systemic, delivery of the drug from the device.

### M1/M2 macrophage ratio

Quadruple staining of pig tissue was performed to reveal the count of M1 and M2 macrophages and to determine the M1/M2 macrophage ratio across all wounds. The antibodies were selected, and the IHC staining protocols were used. The M1 and M2 marker cells were labeled with iNOS and Arginase I, respectively, against the background staining of the pan-macrophage M0 marker BA4D5, as shown in Fig. 7a for control and Flx^+^-treated wounds as representative examples.

**Figure 7.**
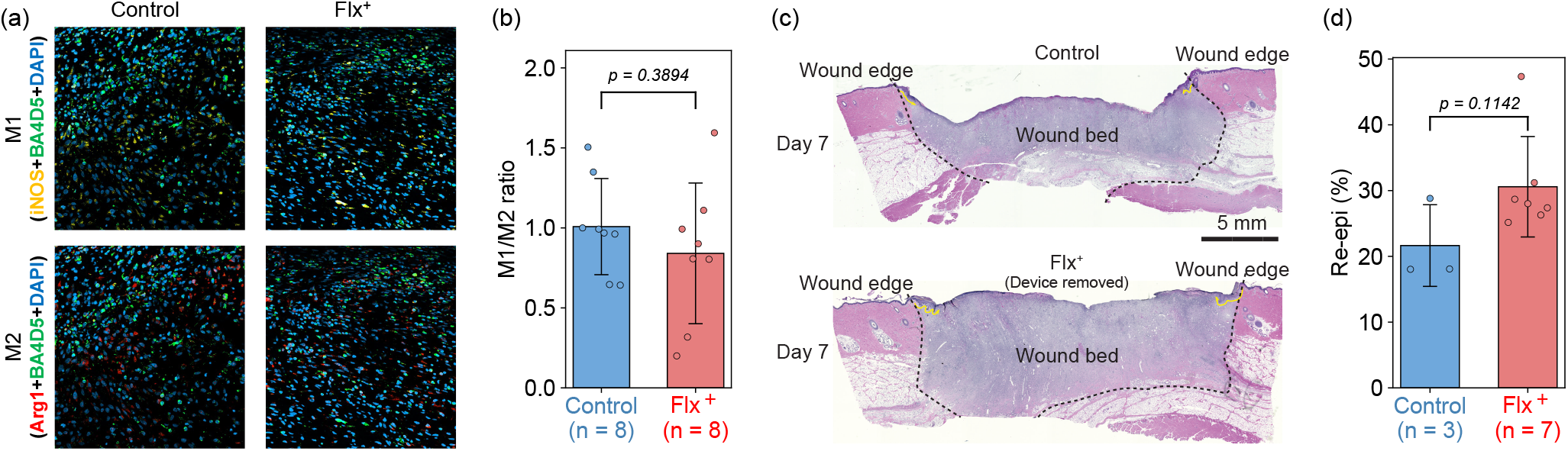
In vivo results for wound tissues harvested on Day 7. (a) Representative IHC images showing M1 macrophages (yellow) and M2 macrophages (red) at the wound edges from control (left) and Flx^+^–treated (right) wounds. (b) Plot of the M1/M2 macrophage ratio for control and Flx^+^–treated wounds. All data presented as mean ± s.d. (c) Representative H&E images showing re-epithelialization for control (top) and Flx^+^-treated (bottom) wounds. The wound edge in each image is outlined by black dotted lines, and re-epithelialization is indicated by yellow arrows extending from the wound edge. (d) Plot of the re-epithelialization percentage for control and Flx^+^–treated wounds. All data presented as mean *±* s.d.

On average, the wound edges of Flx^+^-treated wounds showed a 20% lower M1/M2 macrophage ratio compared to control wounds, based on tissues harvested on Day 7, as shown in Fig. 7b, indicating the advancement of the wound edge earlier into the proliferation (regenerative) phase in the treated wounds. Furthermore, there is a tendency for the M1/M2 macrophage ratio to decrease over time as the wound healing process advances (as seen by comparing Day 3^34^ to Day 7 harvest results in Fig. 7b). From the total number of M1 and M2 cells, the reduction in M1/M2 macrophage ratio at the wound edge appears to be mainly due to the reduction in M1 marker-positive cells. This suggests that Flx^+^ treatment selectively affects M1 macrophages at the wound edge during early stages of wound healing, leading to a shift in the M1/M2 balance toward a less inflammatory state.

### Re-epithelialization

To determine re-epithelialization, 5 *µ*m sections were stained with the H&E staining protocol. H&E images of Flx^+^-treated and control wounds from a 7-day Flx^+^ delivery experiment are shown in Fig. 7c. Re-epithelialization refers to the size of the newly formed epithelium, which is indicated by the yellow arrows. The images were scored to determine the percentage of wound closure (referred to here as re-epithelialization %). Beginning at the innermost edge of the mature collagen (wound edge, outlined by black dotted lines), the lengths of the epithelial tongues and the width of the entire wound (yellow arrows) were measured. From these distances, the percentage of closure for each wound was calculated. As shown in Fig. 7d, the Flx^+^-treated wounds, on average, exhibited 41.67% greater re-epithelialization compared to control wounds, based on tissues harvested on Day 7.

## Summary

In this study, we demonstrate the design, development, and verification of our wireless bioelectronic actuators for wound healing in a porcine model. The devices are used to deliver fluoxetine cations (Flx^+^) and electric field (EF) treatments in full-thickness wounds in pig models. Each treatment can be applied continuously for up to 23 h per day for a period of 7 days. Immunohistochemistry (IHC) staining results show that Flx^+^-treated wounds had an improved M1/M2 macrophage ratio compared to control wounds by 20% at the wound edges. Meanwhile, hematoxylin and eosin (H&E) staining results show that Flx^+^-treated wounds had 41.67% greater re-epithelialization compared to control wounds. More animal experiments are currently being conducted to determine whether statistical significance can be achieved. We are also conducting experiments to better understand and correlate the actuation parameters and biological responses, in order to optimize the timing and dose of the Flx^+^ and EF treatments for accelerating wound healing.

## Methods

### Ethics statement

All pig experiments were conducted under the protocol approved by the University of California Davis (UC Davis) Institutional Animal Care and Use Committee (IACUC). All methods were performed in accordance with the UC Davis IACUC guidelines and regulations. All animal studies are reported in accordance with ARRIVE guidelines.

### PCB fabrication, assembly, and modeling

The CAD model of the PCB is shown in Fig. 1a, which was designed in Autodesk EAGLE. The prototype was fabricated and assembled by a third-party PCB manufacturer. The contact pins were hand-soldered to the plated through holes (PTHs) on the PCB. The 3D model of the PCB was created in Autodesk Fusion 360^35,44^.

### PDMS fabrication, assembly, and modeling

The CAD models of the PDMS ion pump and lid are shown in Fig. 1a, which were created in Autodesk Fusion 360. Each PDMS piece also has its respective 3D mold design (see Supplementary Fig. S10 online). The STL files of the molds were imported into the Formlabs PreForm software. The models were optimized for print orientation, layout, and supports before uploading the resulting file to the Formlabs Form 3 SLA printer for printing with Model V2 resin^35^. Fig. 1a also shows the steps involved in the integration of the PCB with the PDMS device.

### Hydrogel-filled capillary preparation

In order to deliver ions such as Flx^+^, silica capillary tubes are filled with a polyanion hydrogel that selectively conducts cations, following the process detailed by Baniya *et al*.^35^, Dechiraju *et al*.^45^, and Asefifeyzabadi *et al*.^46^. The tubes are then inserted at the bottom of the PDMS device, as shown in Fig. 1a. The hydrogel recipe requires 1 M AMPSA, 0.4 M PEGDA, and 0.05 M photoinitiator (I2959) concentrations to produce a hydrogel with a low swelling ratio (12%) and good conductivity (8.8 ± 0.1 S/m). A several-centimeter length of the silica tubing (inner diameter 0.8 mm, outer diameter 1 mm) used for the capillaries is etched with NaOH and further treated with silane A174 to prevent the hydrogel from swelling out of the capillary after UV crosslinking for 5 min at 8 mW/c*m*^2^. Following UV curing, the capillary tubes are cut into 5 mm pieces. This designated length allows for the easy insertion of capillaries into the PDMS device, with minimal cutting required to reach the final 5 mm length. Cutting the capillary tubing into short lengths beforehand also enables each tube to be checked for electrical connection and loaded with 10 mM Flx HCl solution for Flx^+^ delivery. As shown in Fig. 1a, each device has 9 hydrogel-filled capillary tubes inserted in it.

### Animal experiments

Yorkshire-mixed breed female pigs, weighing 70–80 lbs and approximately 4–5 months old, were used. As illustrated in Supplementary Fig. S1 online, the pigs were acclimated and trained for harness use with positive reinforcement (food treats) for 7–10 days prior to the surgery^47^. Overnight fasting with ad libitum water was applied before both the surgery and device changes. On the surgery day (Day 0), the pigs were induced under general anesthesia, intubated endotracheally, and maintained under anesthesia using masked inhalation of isoflurane (1–5%). Blood samples were collected before surgery and at the endpoint. The back skin was clipped, depilated, and prepared with 2% chlorhexidine scrubs and isopropyl alcohol rinses.

As shown in Fig. 1b, six circular, 20 mm in diameter, full-thickness excisional wounds were created bilaterally on each side of the dorsum in each pig (see Supplementary Fig. S2 online). The four cranial wounds (wounds 1, 2, 5, 6) were treated with the devices. In contrast, the two caudal wounds (wounds 3 and 7) served as standard-of-care controls (no devices applied; wounds were sealed with Conformant 2 wound veil, Optifoam bolsters, and Tegaderm film dressings). The entire wound area was covered with cushion foam sheets and a compression dressing/pig jacket to protect the site. Analgesic medication buprenorphine was administered at extubation, and a fentanyl patch was applied for 72 hours for post-operative pain management.

At the end of the surgery, as illustrated in Fig. 1b, two sets of Raspberry Pi units and power banks—each capable of supporting two devices on the same side—were placed in pouches on each shoulder of the animal and connected to the devices to initiate daily imaging and actuation cycles. Each Raspberry Pi captured images sequentially from its two connected devices, which were applied to two treated wounds, as detailed by Hee *et al*.^42^. The device camera is located above the actuator. As shown in Figs. 1b and 5, the camera consists of a LED ring for brightfield illumination of the wound surface, a lens/sensor stack for image capture, an HDMI connection to the Raspberry Pi for image acquisition/storage, and a USB power connection to the power bank^42^. Wound images were acquired every 2 hours and transmitted wirelessly to a laptop via a local WiFi network for CL control. The imaging cycle lasted the entire 24 hours per day.

Figs. 1b and 5 show the mechanism of EF and/or Flx^+^ delivery from the actuator. A positively charged drug molecule is electrically driven into the wound by applying a positive voltage between the edge and center channels. As a result, ionic currents flow at each channel and can be monitored in real time by the user, as illustrated in Fig. 1b. The actuation cycle lasted between 6 to 23 hours per day until a specific target dose for Flx^+^ (e.g., 0.025 mg per wound) or actuation duration for EF was reached.

The in vivo experiments were conducted over 7 days, with daily wound examination and power bank replacement to initiate new device cycles. Each day began with fully charged power banks to ensure sufficient power for device operation. New imaging and actuation cycles were initiated shortly after plugging in the power banks while the animal stood still with food treats (no additional anesthesia required). At the beginning of Day 3, the devices, wound dressings, cushion foam sheets, and compression jacket were replaced under anesthesia. At the end of the experiment (Day 7 start), the animals were humanely euthanized with pentobarbital > 100 mg/kg IV, the devices were retrieved, and wound tissues were harvested and bisected for IHC and H&E analyses to determine macrophage count and re-epithelialization rate, respectively.

### Fluoxetine in wound tissue and plasma

The wound tissue concentration of fluoxetine was determined by HPLC at the end of topical fluoxetine treatment. Topical fluoxetine was administered daily to pig wounds for 7 days, and wound tissues were collected one day after the final dose was administered. Pig wound tissues (20–30 mg) were cryo-pulverized in liquid nitrogen, resuspended in 0.250 mL methanol containing 0.5% (v/v) formic acid, and sonicated for 15 min in a sonicating bath. The homogenates were centrifuged, and the supernatants were brought to 0.5 mL by adding 0.25 mL water. A 10 *µ*L of the final extract was injected for analysis. Measurements were performed using reverse-phase HPLC with UV absorbance detection. Separation was achieved on an Acquity UPLC BEH C18 column (3.0 mm ID ×100 mm L, 1.7 *µ*m particles) at a flow rate of 0.450 mL/min and a column temperature of 35 °C. The mobile phase consisted of 35:65 acetonitrile:45 mM phosphate buffer (pH = 6.0), and the column effluent was monitored at 230 nm.

To evaluate whether cumulative topical fluoxetine treatment caused systemic exposure, fluoxetine concentrations in pig blood were also measured. Animals received topical fluoxetine on four wounds per day for 7 days, and whole blood samples were collected one day after the final dose was administered. Plasma was separated and fluoxetine extracted and quantified by HPLC. Blood samples were collected in EDTA-treated tubes and centrifuged. Plasma extraction was performed using C18 cartridge solid-phase extraction (SPE) as follows: cartridges were mounted on a vacuum manifold and conditioned with two volumes of methanol and one volume of water. Pig plasma (0.5 mL) was spiked with 50.0 *µ*L internal standard solution (40.0 *µ*g/mL fluvoxamine in methanol), diluted with 0.5 mL H_2_O, and passed through the conditioned SPE cartridges. Cartridges were then rinsed with one volume of H_2_O followed by one volume of 50:50 methanol:H_2_O. Fluoxetine and norfluoxetine were eluted with 0.5 mL methanol containing 0.5% (v/v) formic acid. The eluate was brought to 1.0 mL by adding 0.5 mL H_2_O, and 10 *µ*L was injected for analysis. See Supplementary Fig. S9 online for fluoxetine concentrations in tissue and plasma.

### Histology

The wound tissue was fixed in 4% paraformaldehyde, embedded in paraffin, and sectioned to 5 *µ*m thick for H&E staining to determine the re-epithelialization rate. The H&E stained sections were imaged using a Keyence BZ-9000 inverted microscope under brightfield, and scored with the BZ-II Viewer and Analyzer software (Keyence, Osaka, Japan). The wound edge was defined by the absence of underlying adipose tissue and hair follicles, and the re-epithelialization rate was determined by the percentage of the combined length of the newly formed epidermis^48,49^, as shown previously in Fig. 7c with yellow lines.

### Macrophage IHC staining

To analyze the phenotypic alterations in macrophages during pig wound healing following Flx^+^ and/or EF delivery, we developed a robust IHC protocol featuring quadruple staining to unveil M1/M2 profiles within the same tissue section. This assay encompasses the preparation of experimental pig tissues, involving their embedding in paraffin, followed by sectioning and subsequent staining. This staining process entails the application of specialized antibodies tailored to discern distinct macrophage markers. Within each wound sample, four distinct fluorescent markers are employed, providing the foundation for establishing a comprehensive macrophage panel. These markers encompass: 1) DAPI: Labels all cell nuclei. 2) CD68 (BA4D5): Marks pan-macrophage cells. 3) iNOS: Identifies M1-like macrophages. 4) Arginase I: Identifies M2-like macrophages. Identification of M1 and M2 cells is based on the combination of fluorescent signals. An M1 cell is distinguished by fluorescence in DAPI (blue), CD68 (green), and iNOS (yellow), whereas an M2 cell exhibits fluorescence in DAPI (blue), CD68 (green), and Arginase I (red). This quadruple-staining technique ensures a precise representation for subsequent M1/M2 macrophage ratio calculations. To accomplish this, a customized Cell Profiler pipeline is employed for the recognition of double-positive cells. These cells are subsequently semi-quantified to ascertain the M1/M2 macrophage ratio. This calculation proves pivotal in comparing treated and control wounds, offering vital insights into the efficacy of bioengineered PDMS devices in delivering Flx^+^ and/or EF. This assay plays a critical role in understanding the influence of these treatments on macrophage polarization and, consequently, wound healing.

The IHC protocol is completed over two days. Wound tissue is harvested, fixed in paraformaldehyde, embedded in paraffin and sectioned, and the slides are used for IHC analysis. First the slides are warmed for 30 - 60 min at 60 °C, followed by deparaffinization using 3 xylene washes on a shaker totaling one hour, then followed by 5 min washes in a graded alcohol station, starting from 100% ethanol progressing to 50% ethanol, and then finally ending with PBS washes. Slides then undergo the antigen retrieval step in a rice cooker/vegetable steamer for 40 min, in a bath of EDTA Ph 8.0. Slides are cooled for 30 min. PBS/PBST washes are performed for 1 h total. Slides are then blocked with 10% Donkey Serum + 3% BSA + PBST (tritonx-100 at 1%) for 2 hours at room temperature in a humidified slide box. Primary antibodies are then incubated overnight at room temperature for 12-16 h in a humidified slide box. Primary antibodies are incubated at the following dilutions: iNOS 1:50 (Thermo fisher, cat # PA1036), Arginase I 1:50 (Thermo fisher, cat # PA518392), and BA4D5 1:20 (Bio-Rad, cat # MCA2317GA). The next day, slides are washed for 1 hour in PBS and PBST. Then incubated for 1.5 h in 1:200 diluted secondary antibodies dyed with different Alexafluors. Afterward, a TRUEVIEW autofluorescence kit is added for 5 min, followed by a 15 min incubation in DAPI. Then slides are mounted with nail polish and imaged on Zeiss Confocal LSM 900. After imaging, photos are viewed through ImageJ and analyzed using a program called Cell Profiler to label colocalization and assist in manual counting of each image. Eight images are taken, 4 in the wound center and 2 at both wound edges.

### PDMS design

The PDMS device is inserted into the wound for Flx^+^ and/or EF delivery, as shown in Fig. 1b. The design and fabrication of the PDMS device follow a process similar to that outlined by Baniya *et al*.^35^. Fig. 1a shows the CAD model of the PDMS device, which consists of two pieces: the ion pump and the lid, each of which is fabricated separately from its respective 3D-printed mold (see Supplementary Fig. S10 online). The molds are filled with PDMS and then cured. The two PDMS pieces are then demolded and bonded together. A clamp is used to bond the two pieces to ensure good manufacturing yield.

The PDMS ion pump contains reservoirs to store the drug solution of interest and through-holes for electronic interfacing. The PDMS lid seals the reservoirs and contains a notch with a diameter of 20 mm and a depth of 8–10 mm, which roughly matches the wound dimensions. The notch has nine vertical channels (eight at the edge, one at the center) that connect the reservoirs to the wound bed. Hydrogel-filled capillaries are inserted at the bottom of the channels to prevent solution leakage while allowing the passage of positively charged ions/biomolecules, such as Flx^+^, to the wound upon voltage application.

The PDMS lid thickness (optical path) is chosen so that the camera can maintain a clear focus on the wound bed through its lens, as shown in Figs. 1b and 5. A deep cut is made at the wounding site to remove the thick fat layer, allowing the notch to be inserted and reach the muscle layer, as shown in Figs. 1b and 5.

## Supplementary information

See supplementary information for Figs. S1 to S10 and the supplementary text.

## Acknowledgements

This research is supported by the Defense Advanced Research Projects Agency (DARPA) through Cooperative Agreement Number D20AC00003 awarded by the U.S. Department of the Interior (DOI), Interior Business Center.

## Author contributions

P.B. designed, fabricated, and characterized the PCBs; developed the device GUI, software, firmware, and closed-loop system; conceptualized the PCB-PDMS integration, and wrote the main manuscript. M.Tebyani, K.S., and A.B. designed and fabricated the PDMS and integrated the PDMS with the PCB. W.S.H. and G.K. designed and integrated the camera. H.L., N.A., C.H., T.N., and K.D. prepared the capillary and performed ion-pump experiments. H.Y. conducted the animal experiments. A.G. performed the H&E staining and quantified re-epithelialization. K.Z. and C.R. performed the IHC staining and quantification of M1/M2 macrophage ratio. F.L. wrote the wound healing algorithm. C.F. 3D-printed the device enclosures. E.A. coordinated the experiments. M.R., M.Teodorescu, M.G., M.Z., and R.R.I. secured the funding and directed the research. All authors reviewed the manuscript.

## Data availability

The data generated and analyzed during this study are available from the corresponding author on reasonable request.

