## Supplementary Information for "Development of a wireless bioelectronic actuator for wound healing in a porcine model"

### Supplementary figures and text

#### In vivo experimental procedures and setup

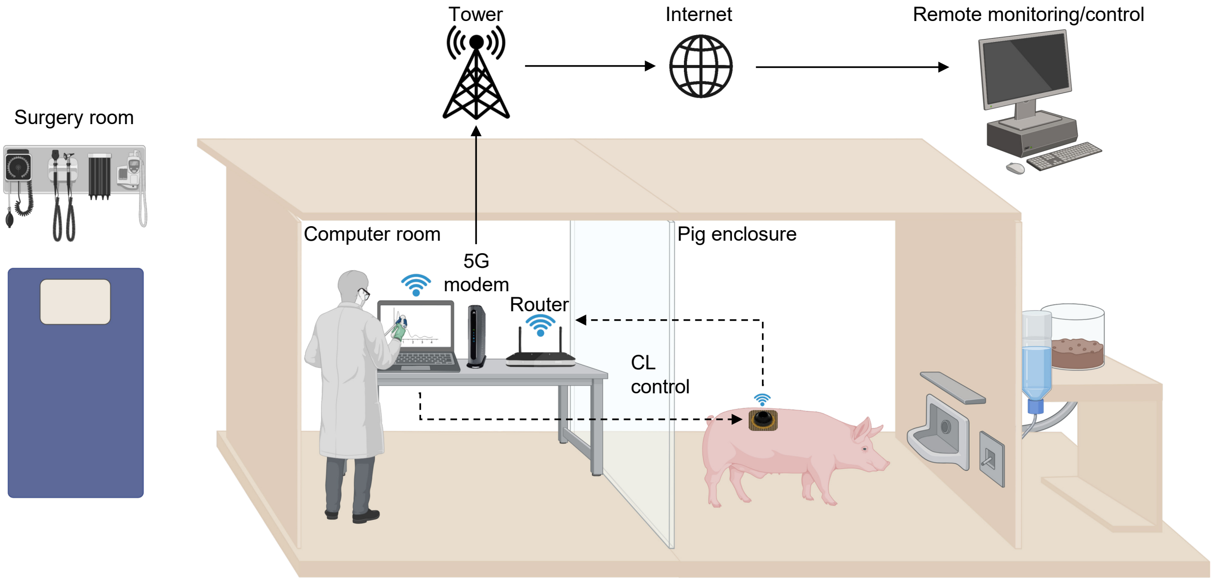


**Figure S1.** The pig is first anesthetized in a surgery room where wounds are created, and the wireless bioelectronic devices are applied. The pig is then transferred to a nearby enclosure. After that, closed-loop (CL) control of $Flx^{+}$ and/or EF delivery can be initiated, and the device transmits real-time data to the laptop via WiFi in an adjacent room operated by a researcher. The setup also supports remote monitoring and control through 5G internet connectivity, enabling real-time oversight by researchers. The animal has access to water and food, ensuring its welfare during long-term monitoring studies.

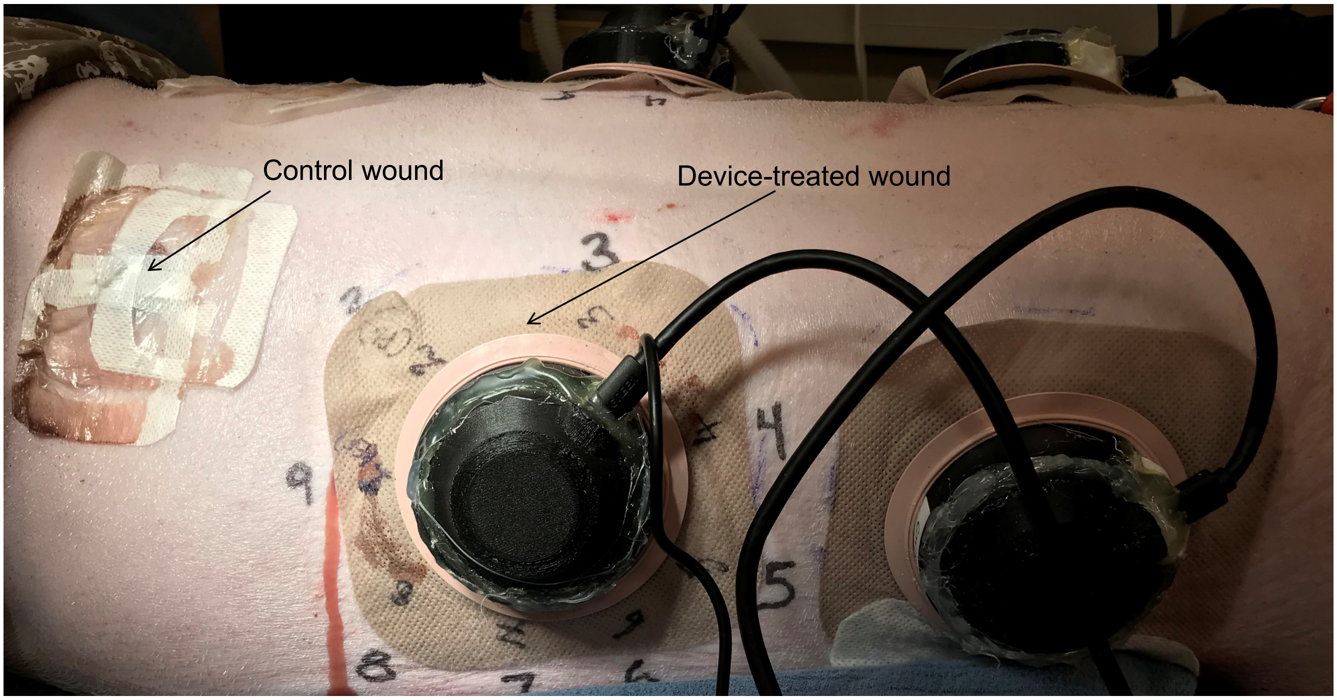


**Figure S2.** Device-treated and control (standard-of-care) wounds on the back of a pig. Wireless bioelectronic devices are secured to the treated wounds with the help of a skin barrier and tape.

### ADC and I-DAC calibration

The ideal (uncalibrated) output code $A_i$ of the ADC at each channel $i$, or electrode $E_i$, for an input voltage $V_i$, or electrode voltage $V_{E_i}$, is given by

$$A_{i/E_i} = \frac{2^{N_{adc}}}{V_{CC}} V_{i/E_i} \tag{1}$$

Two-point calibration of the ADC offset and gain is performed to enable accurate voltage and current measurements at each channel. Input voltages chosen for calibration are $V_c^1 = 0$ V and $V_c^2 \approx 4.7$ V, which are set by putting the I-DAC in voltage output mode and setting its output to the minimum and maximum, respectively. These values were chosen as calibration points because they fall within the linear range of the ADC. All channels must be left open-circuited during voltage calibration, during which ADC measurements $A_{c_i}^1$ and $A_{c_i}^2$ for the two calibration points are made at each channel $i$ by the device internally. The corresponding voltage measurements $V_{c_i}^1$ and $V_{c_i}^2$ for the two calibration points are made at each channel $i$ externally using a multimeter, as shown in Fig. S3, but without the resistor. The calibrated output code $A_{c_i}$ of the ADC at each channel $i$ is given by

$$A_{c_i} = m_{adc_i} V_{c_i} + b_{adc_i} \tag{2}$$

where

$$m_{adc_i} = \frac{A_{c_i}^2 - A_{c_i}^1}{V_{c_i}^2 - V_{c_i}^1} \tag{3}$$

and

$$b_{adc_i} = A_{c_i}^1 - m_{adc_i} V_{c_i}^1 \tag{4}$$

are the ADC calibration coefficients.

After calibration, the resistor is put back in series at each channel individually, as shown in Fig. S3, and the I-DAC output code is swept from 0 to 255 to vary the ADC input voltage from 0 to 4.7 V. The ideal, calibrated, and measured ADC voltage transfer functions for all channels are shown in Fig. S4.

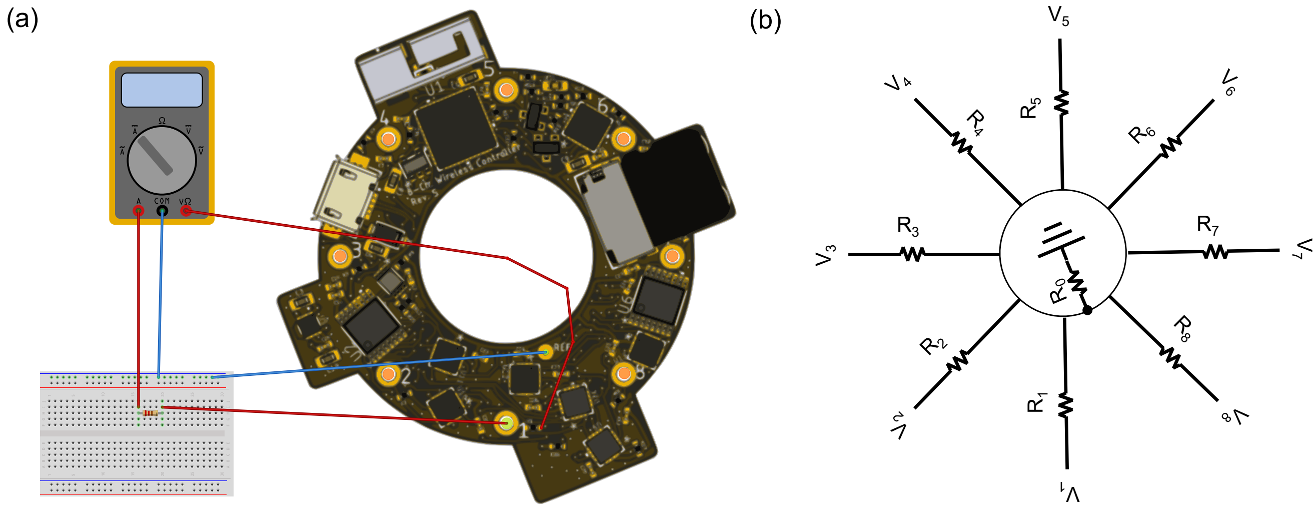


**Figure S3.** (a) To calibrate the ADCs and DACs, a multimeter and a breadboard are used to measure the current flowing through and the voltage across a 46.13 kΩ resistor at each channel (PTH) of the PCB. (b) A circuit diagram for in-house PCB testing, where each channel is loaded with resistors to emulate a device delivering drugs to the wound.

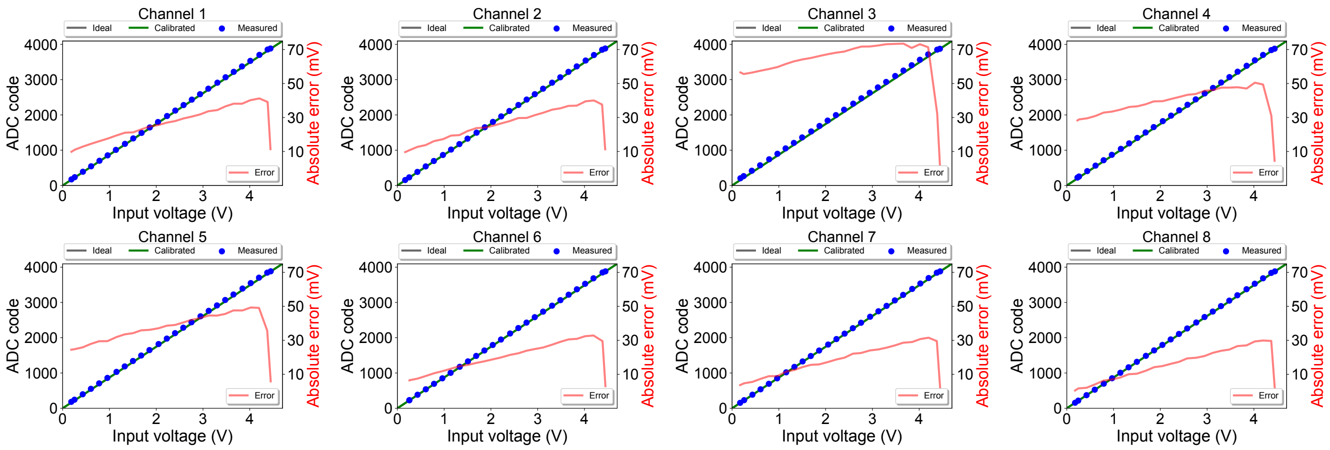


**Figure S4.** ADC voltage transfer functions and absolute voltage errors for all 8 channels.

The ideal (uncalibrated) output current $I_i$ of the I-DAC at each channel $i$, for an input code $D_i$, is given by

$$I_i = \frac{I_{max}}{2^{N_{idac}}} D_i \tag{5}$$

Two-point calibration of the I-DAC offset and gain is performed to enable accurate current measurements at each channel. Nominal currents chosen for calibration are designated as $I_c^1 \approx 2\ \mu$A and $I_c^2 \approx 50\ \mu$A, corresponding to I-DAC codes $D_c^1 = 1$ and $D_c^2 = 255$, respectively. These values were chosen as calibration points because they fall within the linear range of the I-DAC. Each channel has a 46.13 kΩ calibration resistor connected between it and ground (center channel, 0 V) to allow current flow during calibration, during which multimeter measurements $I_{c_i}^1$ and $I_{c_i}^2$ for the two calibration points are made at each channel $i$. These currents are measured using a multimeter, as shown in Fig. S3. The calibrated output current $I_{c_i}$ of the I-DAC at each channel $i$ is given by

$$I_{c_i} = m_{idac_i} D_i + b_{idac} \tag{6}$$

where

$$m_{idac_i} = \frac{I_{c_i}^2 - I_{c_i}^1}{D_{c_i}^2 - D_{c_i}^1} \tag{7}$$

and

$$b_{idac_i} = I_{c_i}^1 - m_{idac_i} D_{c_i}^1 \tag{8}$$

are the I-DAC calibration coefficients.

The I-DAC output code is swept from 0 to 255 to vary the I-DAC output current from 0 to 50 $\mu$A.. The ideal, calibrated, and measured DAC transfer functions for all channels are shown in Fig. S5.

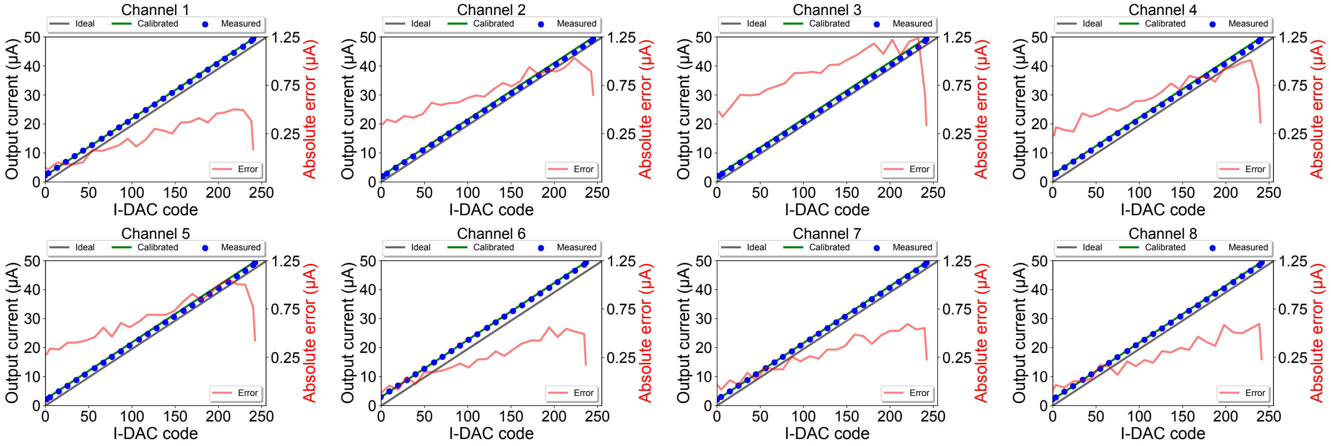


**Figure S5.** I-DAC current transfer functions and absolute current errors for all 8 channels.

The difference (diff.) of two uncalibrated ADC codes, $A_i$ at channel $i$ and $A_{E_i}$ at electrode $i$, is related to the input current $I_i$ at channel $i$, as follows:

$$A_i - A_{E_i} = I_i(\mu\text{A})\,0.01\,\frac{2^{N_{adc}}}{V_{CC}} \tag{9}$$

where $I_i$ is the input current calculated from uncalibrated input voltages, ranging from 0 to 50 $\mu$A (as I-DAC code $D_i$ is varied from 0 to $2^{N_{dac}} - 1$), and is given by

$$I_i(\mu\text{A}) = (V_i - V_{E_i})\,100 \tag{10}$$

The difference (diff.) of two calibrated ADC codes, $A_{c_i}$ at channel $i$ and $A_{E_{c_i}}$ at electrode $i$, is related to the current $I_i$ at channel $i$, as follows:

$$A_{c_i} - A_{E_{c_i}} = I_{c_i}(\mu\text{A})\,0.01\,m_{adc_i} + b_{adc_i} - b_{adc_{E_i}}\frac{m_{adc_i}}{m_{adc_{E_i}}} \tag{11}$$

where $I_{c_i}$ is the input current calculated from calibrated input voltages, ranging from 0 to 50 $\mu$A (as the I-DAC code $D_i$ is varied from 0 to $2^{N_{dac}}-1$), and is given by

$$I_{c_i}(\mu\text{A}) = (V_{c_i} - V_{c_{E_i}})\,100 \tag{12}$$

The I-DAC code is swept from 0 to 255 to vary the ADC input current from 0 to 50 $\mu$A. The ideal, calibrated, and measured ADC current transfer functions for all channels are shown in Fig. S6.

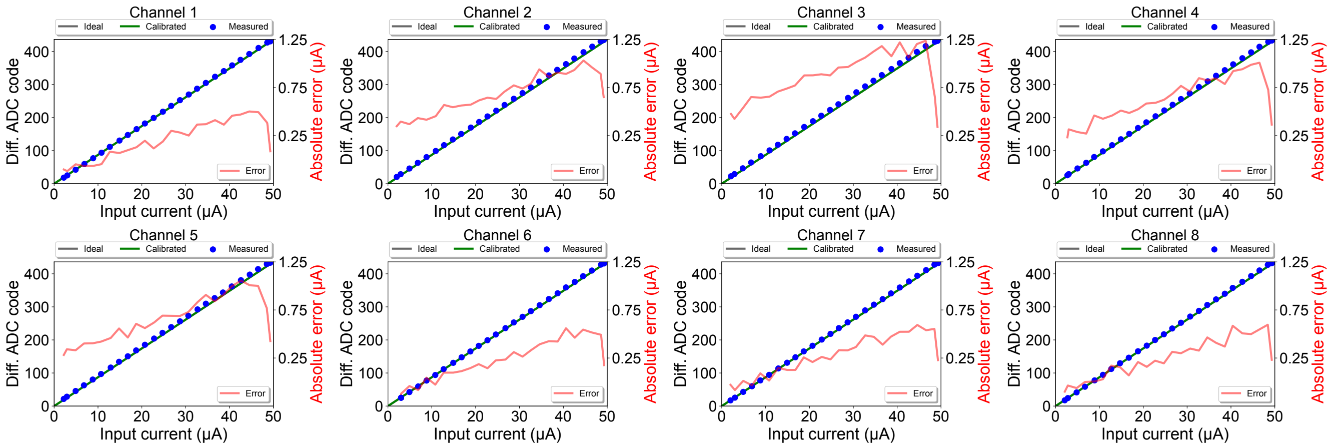


**Figure S6.** ADC current transfer functions and absolute current errors for all 8 channels.

### Graphical user interface (GUI)

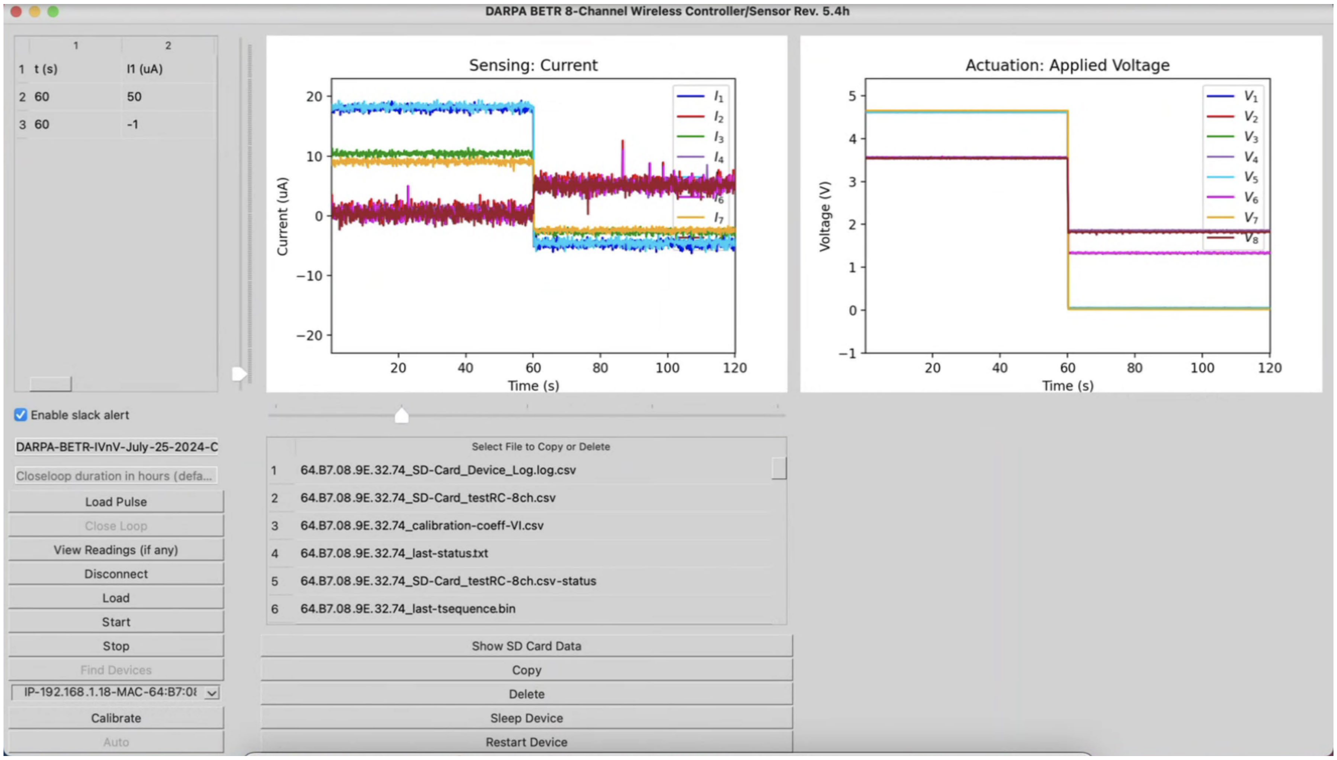


**Figure S7.** Graphical user interface (GUI) for data acquisition and control of wireless bioelectronic actuators. The GUI supports simultaneous operation of multiple devices.

### Antenna

Fig. S8a also shows how the antenna and its 3D radiation pattern are aligned. The pattern is broader, and the gain is higher along the positive x-axis (the right side of the antenna), as evident in the horizontal (xy) plane pattern in Fig. S8b. The vertical plane patterns show that the radiation above (+z-axis) and below (-z-axis) the antenna is approximately the same. The vertical (yz) plane pattern indicates that the gain is higher along the positive y-axis (the front of the antenna).

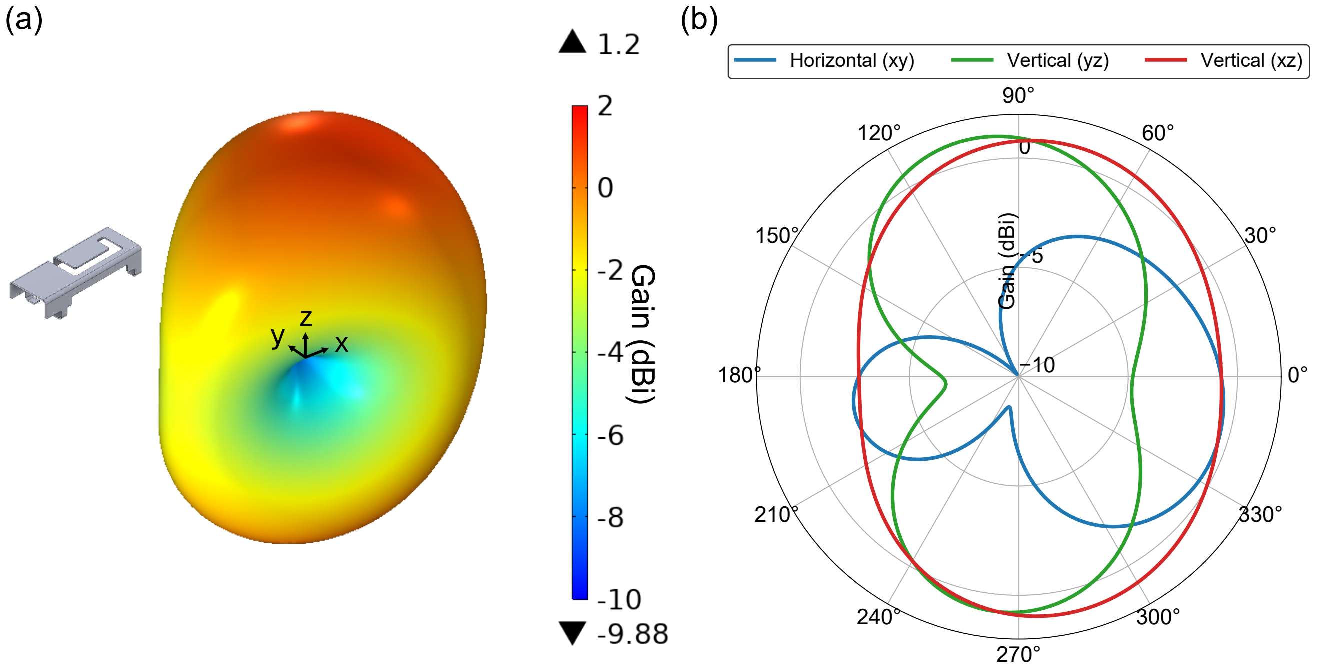


**Figure S8.** Simulated radiation pattern of the 3D metal antenna on the PCB at 2.45 GHz. (a) 3D gain pattern. (b) 2D gain patterns in the horizontal (xy) and vertical (yz, xz) planes.

#### Fluoxetine concentration

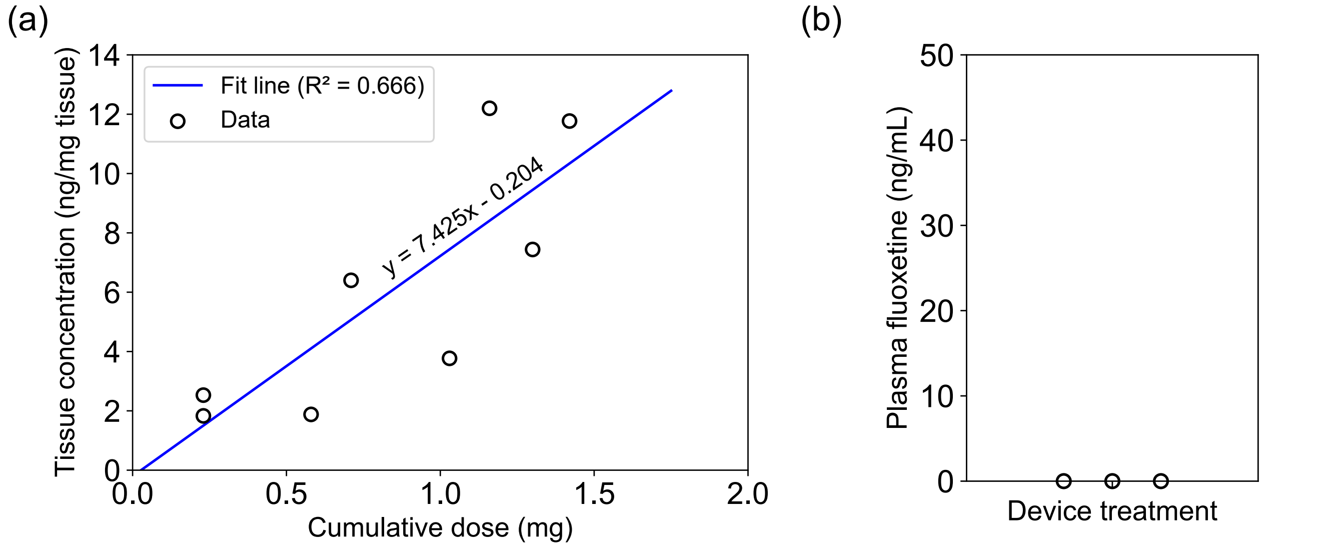


**Figure S9.** (a) The wound tissue concentration of fluoxetine was strongly correlated with the cumulative dose delivered by the experimental device (Spearman's r = 0.862, p = 0.005873, from n = 8 wounds treated with $Flx^+$ devices from 3 independent experiments). The cumulative dose represents the total amount of fluoxetine delivered to each wound by the device over the course of the 7-day treatment. (b) Fluoxetine was not detectable in pig plasma (n = 3 pigs) following topical fluoxetine wound treatment using the experimental device.

**PDMS fabrication**

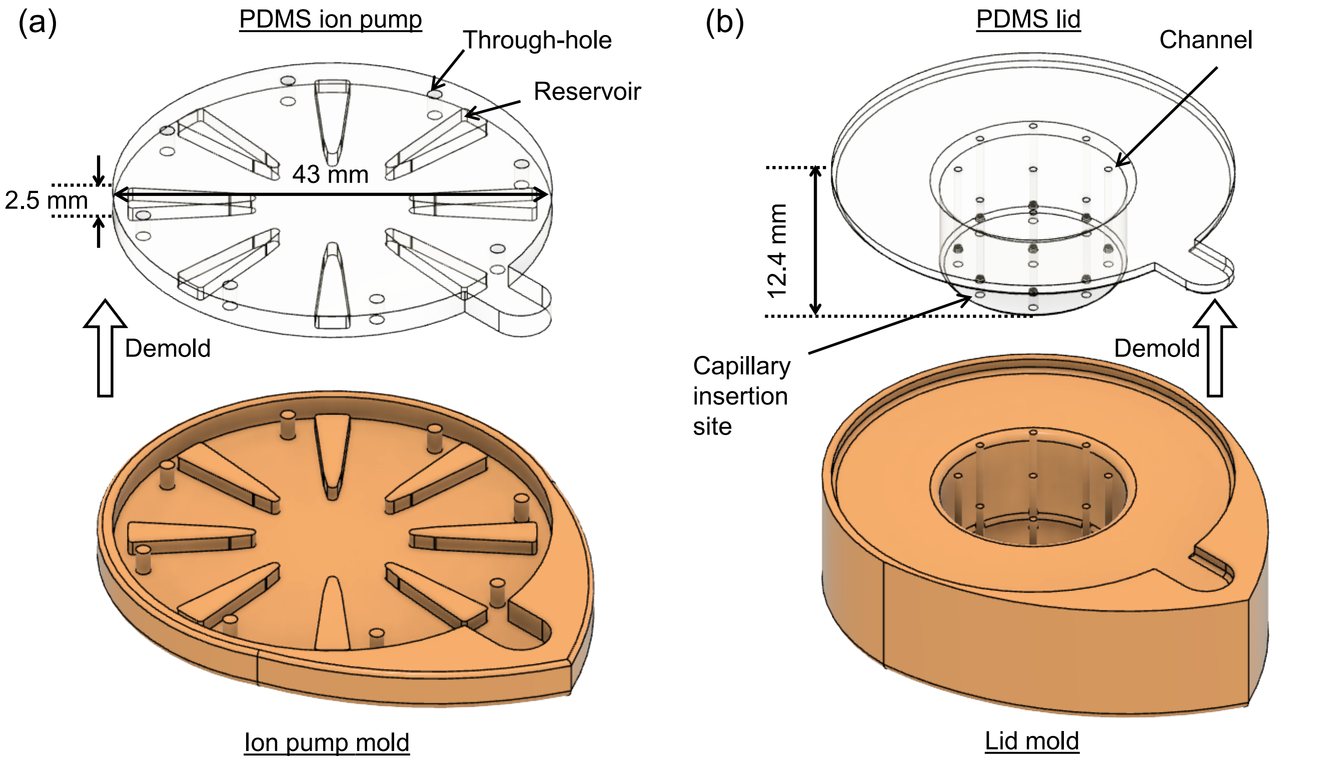


**Figure S10.** Fabrication process of the PDMS device using 3D-printed two-part molds. (a) PDMS ion pump and its mold. (b) PDMS lid and its mold.
